# Role of α-Catenin and its mechanosensing properties in the regulation of Hippo/YAP-dependent tissue growth

**DOI:** 10.1101/567750

**Authors:** Ritu Sarpal, Victoria Yan, Lidia Kazakova, Luka Sheppard, Ulrich Tepass

## Abstract

α-catenin is a key protein of adherens junctions (AJs) with mechanosensory properties. It also acts as a tumor suppressor that limits tissue growth. Here we analyzed the function of Drosophila α-Catenin (α-Cat) in growth regulation of the wing epithelium. We found that different α-Cat levels led to a differential activation of Hippo/Yorkie or JNK signaling causing tissue overgrowth or degeneration, respectively. α-Cat can modulate Yorkie-dependent tissue growth through recruitment of Ajuba, a negative regulator of Hippo signaling, to AJs but also through a mechanism that does not involve junctional recruitment of Ajuba. Further, both mechanosensory regions of α-Cat, the M region and the actin-binding domain (ABD), contribute to growth regulation. Whereas M is dispensable for α-Cat function in the wing, individual M domains (M1, M2, M3) have opposing effects on growth regulation. In particular, M1 limits Ajuba recruitment. Loss of M1 cause Ajuba hyper-recruitment to AJs promoting tissue-tension independent overgrowth. Although M1 binds Vinculin, Vinculin it is not responsible for this effect. Moreover, disruption of mechanosensing of the α-Cat actin-binding domain affects tissue growth, with enhanced actin interactions stabilizing junctions and leading to tissue overgrowth. Together, our findings indicate that α-Cat acts through multiple mechanisms to control tissue growth, including regulation of AJ stability, mechanosensitive Ajuba recruitment, and dynamic direct F-actin interactions.

## Introduction

The cadherin-catenin complex (CCC) at adherens junctions (AJs) links actomyosin networks of neighboring cells thereby sensing and distributing cytoskeletal tension across tissues [Takeichi, 2014, 2018]. α-catenin physically couples the cadherin-β-catenin complex to the actin cytoskeleton. Loss of α-catenin function, similar to the loss of E-cadherin, disrupts epithelial integrity including a loss of apical-basal polarity, cell adhesion, and the ability of cells to undergo coordinated cell movements. α-catenin can also act as a mechanosensor. Biochemical responses to mechanosensing are thought to have multiple consequences [Pinhero and Bellaiche, 2018; Charras and Yap, 2018; Yap et al., 2018], including a strengthening of F-actin binding to enhance cell adhesion [Yonemura et al., 2010; Buckley et al., 2014; Ishiyama et al., 2013; 2018], reorganization of actin at cell junctions [Ishiyama et al., 2018], and modulation of cell signalling that regulates tissue growth through the Hippo/YAP pathway [Rauskolb et al., 2014]. However, in vivo evidence for specific functions of α-catenin-mediated mechanosensing remains very limited.

E-cadherin and α-catenin act as tumor suppressor proteins [Birchmeier and Behrens, 1994; Benjamin and Nelson, 2008; Vasioukhin, 2012]. Deregulation of the transcriptional co-activator YAP (Yorkie [Yki] in Drosophila) is one mechanism of how the loss of the CCC can impact tissue growth [Karaman and Halder, 2018]. YAP is the key effector of the Hippo-Warts kinase cascade (MST1/2 and LATS1/2 in mammals) [Misra and Irvine, 2018]. α-catenin can regulate YAP/Yki through multiple, potentially cell type-specific mechanisms including cytoplasmic sequestration or activation of an integrin-Src pathway [Silvis et al., 2011; Schlegelmilch et al., 2011; Li et al., 2016]. Moreover, active Yki not only acts in the nucleus to stimulate growth but is also recruited to apical junctions where it enhances myosin II activity, which in turn promotes tissue growth through the Hippo/Yki pathway [Xu et al., 2018].

It was proposed recently that mechanosensing of Drosophila α-Catenin (α-Cat) can modulate the Hippo/Yki pathway via the LIM domain protein Ajuba (Jub), a negative regulator of Warts (Wts). In response to cytoskeletal tension, α-Cat sequesters a Jub-Wts complex at AJs, preventing the phosphorylation of Yki and causing tissue growth [Das Thakur et al., 2010; Rauskolb et al., 2014; Pan et al., 2016]. A similar AJ-dependent mechanism appears to be conserved in mammals [Karaman and Halder, 2018; Misra and Irvine, 2018]. However, loss of function of α-Cat, DE-cadherin (DEcad), and the Drosophila β-catenin protein Armadillo (Arm) were reported to have a negative impact on growth and caused reduced Yki activity in mutant cells and enhanced JNK signaling leading to cell death [Yang et al., 2015]. Here, we address these discrepancies and found that depending on the degree of α-Cat loss a differential activation of the Hippo/Yki and JNK pathways is observed, which causes either tissue overgrowth driven by Yki activation or tissue undergrowth driven by JNK elicited cell death.

One key feature of α-catenin is its mechanosensory properties. The α-catenin protein comprises three evolutionary conserved regions: a N-terminal region that binds to β-catenin, a central M-region, and a C-terminal actin-binding domain (ABD). The activity of cytoplasmic myosin II applies tension to α-catenin when bound to β-catenin and F-actin, causing at least two conformational changes: First, the N-terminal α-helix within the ABD (α1-helix) is pulled away from ABD revealing an enhanced F-actin binding interface and facilitating dimerization of ABD to promote actin bundling [Ishiyama et al., 2018]. This mechanosensory property of α-catenin ABD explains its catch bond behavior, the ability of a chemical bond to strengthen as a result of force application [Buckley et al., 2014]. Second, the M-region of α-catenin also undergoes a conformational change when stretched. M consists of three α-helical bundles, the M1, M2, and M3 domains. The angle between the M2 and M3 domains increases upon force application and the M1 domain unfurls to reveal a cryptic binding site for the actin-binding protein Vinculin [Yonemura et al., 2010; le Duc et al., 2010; Choi et al., 2012; Ishiyama et al., 2013; Yao et al., 2014]. The recruitment of Vinculin is thought to strengthen the AJ-actin link. However, in vivo evidence supporting this hypothesis is limited.

The Drosophila wing disc epithelium has been a crucial model to investigate the regulation of tissue growth [Hariharan, 2015]. The JNK and Hippo/Yki pathways monitor epithelial health and play key roles in the regulation of disc size. Tissue growth in the wing epithelium is also influenced by mechanical tension [Eder et al., 2017]. Differences in tension correlate with Yki activation, a response thought to be mediated by AJ mechanosensing and the Hippo pathway [Rauskolb et al., 2014; Pan et al., 2016; 2018]. We have tested this model by replacing endogenous α-Cat with mutant forms that compromise α-Cat M-region or ABD-based mechanosensing, and asked how these α-Cat mutant isoforms can affect AJ stability, Jub recruitment and tissue growth.

## Results

### α-Catenin function in growth regulation and epithelial maintenance can be genetically separated

Our previous work suggested that the levels of the CCC can be reduced to a few percent of normal and still maintain normal AJs and epithelial integrity in tissues that do not undergo cell rearrangements such as the late embryonic epidermis [Tepass et al., 1996; Sarpal et al., 2012]. To test whether α-Cat has a cell-autonomous function in growth regulation we took advantage of the wing disc (Figure 1A), which shows dramatic tissue growth but little cell rearrangement during larval development. Based on findings in the embryo, we reasoned that a moderate reduction of α-Cat would not disrupt epithelial polarity causing cell death in the developing wing, and hence could reveal defects in tissue growth. We examined tissue in which α-Cat was reduced to different degrees (a phenotypic series) starting with the null phenotype. As *α-Cat* mutants are embryonic lethal [Sarpal et al., 2012] we examined cell clones mutant for a null allele of *α-Cat*. In contrast to control clones, we did not detect *α-Cat* null clones in the third larval wing imaginal disc (Figure 1A,B) [Sarpal et al., 2012]. We expected that the loss of AJs initiates programmed cell death [Kolahgar et al., 2011; Yang et al., 2015]. Upon expression of the caspase inhibitor p35 [Hay et al., 1994] we found that *α-Cat* null cells accumulate below the disc epithelium and display spindle-shaped morphologies with extensive protrusions (Figure 1B). We conclude that wing epithelial cells devoid of α-Cat undergo programmed cell death, and if prevented from dying leave the epithelium and adopt a mesenchyme-like character.

**Figure 1:**
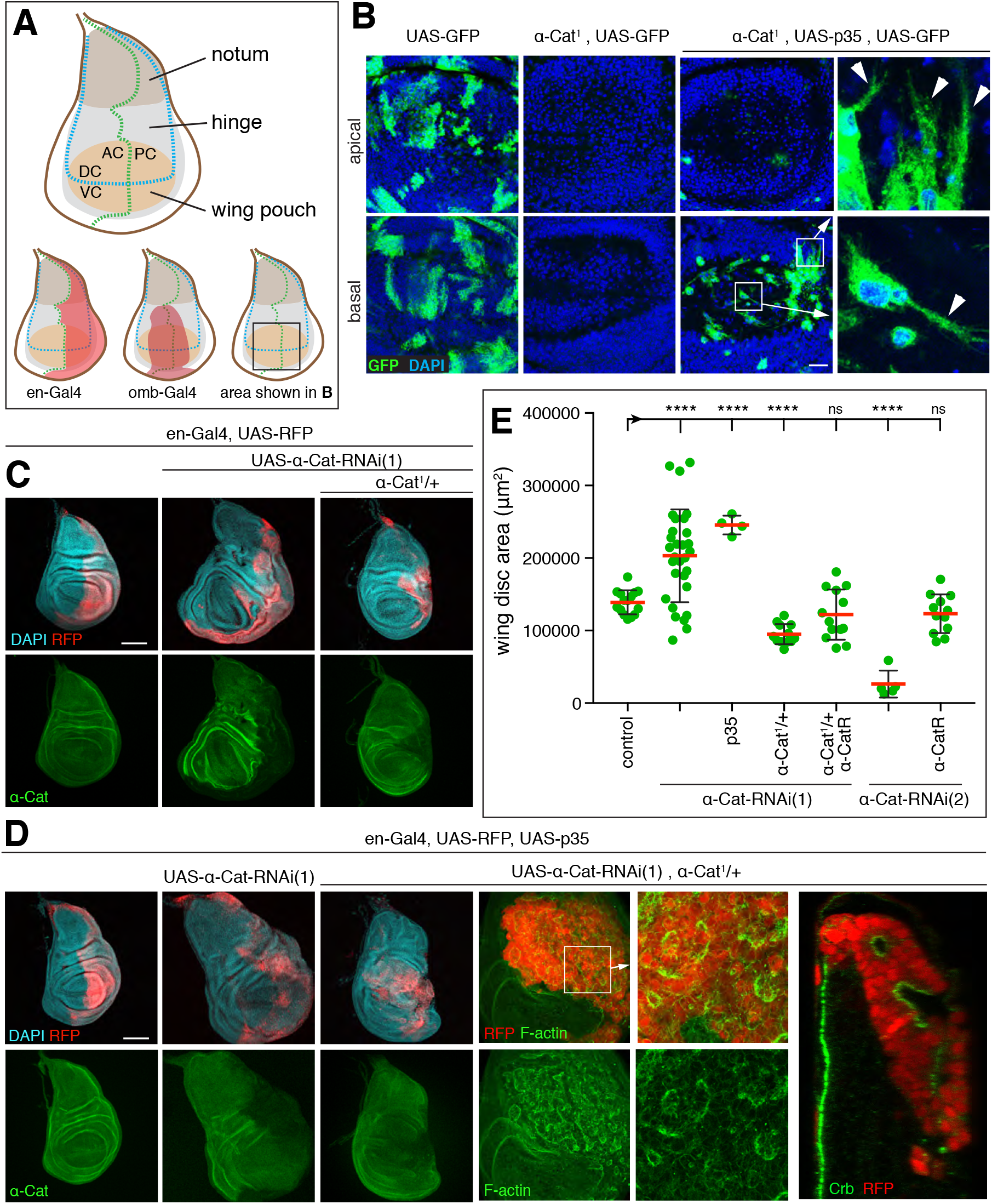
A phenotypic series for α-Cat reveals distinct roles in growth regulation and epithelial polarity. (**A**) Schematic of 3^rd^ larval instar wing imaginal disc. Indicated are the main subdivisions of the disc proper, the compartment boundaries (AC, anterior compartment; PC posterior compartment; DC, dorsal compartment; VC, ventral compartment), the expression domains of the *en-Gal4* and *omb-Gal4* drivers used in this study, and the area of the discs shown in (B). (**B**) Late 3^rd^ larval instar wing discs with control (left two panels) or *α-Cat^1^* null mutant clones positively labeled with GFP. In contrast to control clones, *α-Cat^1^* mutant clones are not observed. When cell death is suppressed through expression of p35, *α-Cat^1^* mutant cells are found basal to the epithelium and show extensive protrusive activity (arrowheads in close-ups). Scale bar, 25 μm. (**C**) KD of α-Cat in the PC (marked by RFP) with *α-CatRNAi(1)* causes tissue overgrowth, whereas KD of α-Cat in the PC with *α-CatRNAi(1)* in the presence of one copy of *α-Cat^1^* causes a degeneration of the PC. Scale bar, 100 μm. (**D**) KD of α-Cat in the PC (marked by RFP) with *α-CatRNAi(1)* while expressing p35 causes tissue overgrowth, whereas KD of α-Cat in the PC with *α-CatRNAi(1)* while expressing p35 in the presence of one copy of *α-Cat^1^* causes the formation of a multilayered tumor mass with small epithelial vesicles or patches. Apical domain of epithelial cells marked by enrichment of F-actin and Crb. Scale bars, 100 μm. (**E**) Quantification of wing disc area in flies of indicated genotypes. Two-tailed, unpaired t-test used to determine statistical significance. ns (P>0.05), ****(P<0.0001).

To elicit a less drastic reduction of α-Cat function we expressed two different *α-Cat* shRNAs in the posterior compartment (PC) of the wing disc with the *en-Gal4* driver (Figure 1A). *α-Cat-RNAi(1)* is directed against the 5’UTR of *α-Cat* whereas *α-Cat-RNAi(2)* targets a region in the *α-Cat* RNA that encodes the M2 domain. *en>α-Cat-RNAi(1)* caused hyperplastic overgrowth of the wing disc with both enlarged anterior compartment (AC) and PC suggesting non-cell-autonomous and cell-autonomous tissue overgrowth (Figure 1C,E). Further reduction of α-Cat by expression of *α-Cat-RNAi(1)* in animals that carried one mutant copy of *α-Cat* (*en>α-Cat-RNAi(1), α-Cat/+*) caused a reduction in disc size due to a degeneration of much of the PC (Figure 1C,E). An even more pronounced degeneration of the PC was seen upon expression of *α-Cat-RNAi(2)* (Figure 1E, and below). This indicates that *α-Cat-RNAi(2)* causes a stronger knockdown (KD) of α-Cat than *α-Cat-RNAi(1)*. Moreover, whereas the degeneration of the PC upon α-Cat KD has been reported previously [Yang et al., 2015], we found that a moderate KD of α-Cat caused a conspicuous tissue overgrowth, suggesting that α-Cat normally limits tissue growth.

We combined *α-Cat* KD with p35 expression to examine how cell death contributes to the KD phenotypes. *en>α-Cat-RNAi(1) p35* discs were larger compared to *en>α-Cat-RNAi(1)* discs (Figure 1D,E), suggesting that the hyperplastic discs resulting from moderate *α-Cat* KD showed a significant amount of cell death. Suppression of cell death in strong *α-Cat* KD conditions (*en>α-Cat-RNAi(1) p35, α-Cat/+*) led to cell-autonomous formation of multilayered tissue masses (Figure 1D). These tumor-like masses contained pockets of epithelial cells as was evident from labeling with the apical markers Crumbs (Crb) and F-actin (Figure 1D). Taken together, our analysis of *α-Cat* loss-of-function conditions in conjunction with a block of cell death defined three distinct phenotypic classes that align with the progression sequence of epithelial cancer from adenoma (epithelial overgrowth), to adenocarcinoma (overgrowth associated with a partial loss of epithelial integrity), to carcinoma (loss of epithelial integrity with cells showing protrusive activity).

### Differential activation of JNK and Hippo/Yorkie signaling in α-Cat compromised wing epithelia

Activation of JNK signaling that causes cell death under strong *α-Cat* KD conditions was reported previously (Yang et al., 2015). We confirmed these data (not shown). In addition, we tested whether JNK is activated with moderate *α-Cat* KD. Using the JNK transcriptional reporter *puc^E697^-lacZ* (*puc-lacZ*) we found that the JNK pathway was activated in *en>α-Cat-RNAi(1)* and *en>α-Cat-RNAi(1) p35* discs, in which cell death is blocked downstream of JNK activation through p35 expression (Figure 2A). Interestingly, *α-Cat-RNAi(1) puc-lacZ* discs did not show hyperplastic growth as observed for *α-Cat-RNAi(1)* but a strong degeneration of the PC. *puc-lacZ* is a loss of function allele of *puckered (puc)* which should lead to higher JNK activity levels as Puc is a phosphatase that negatively regulates JNK signaling through dephosphorylation of the JNK kinase Basket (Figure 2B) [Riesgo-Escovar et al., 1996; Martin-Blanco et al., 1998]. The genetic interaction between *α-Cat-RNAi(1)* and *puc-lacZ* is consistent with the notion that JNK signaling is activated in *α-Cat-RNAi(1)* discs. However, the levels of JNK-elicited cell death in *en>α-Cat-RNAi(1)* discs are apparently insufficient to overcome the concomitant tissue overgrowth resulting from the depletion of α-Cat.

**Figure 2:**
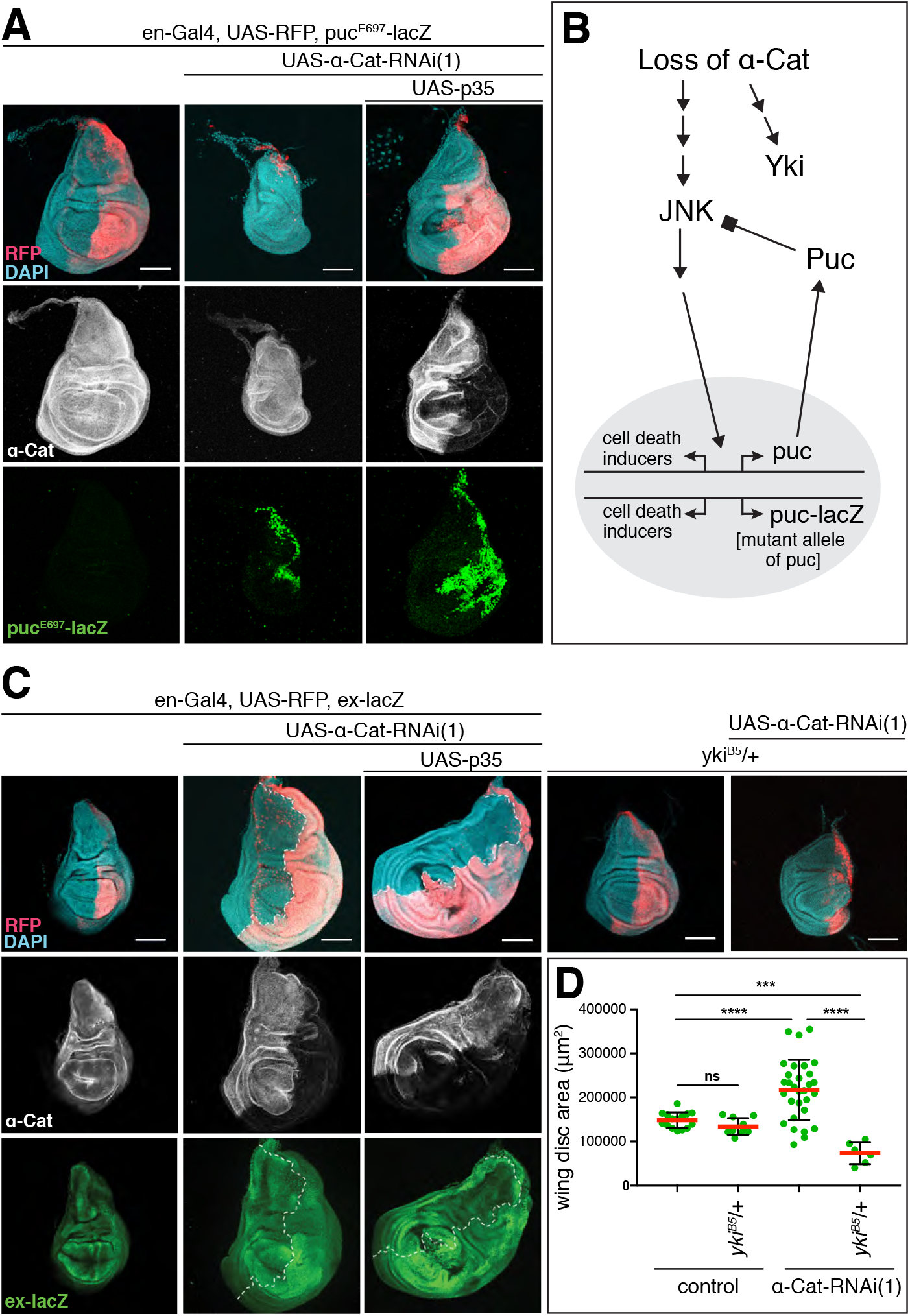
Loss of α-Cat activates JNK pathway and Yki. (**A**) Activation of the JNK transcriptional reporter *puc-LacZ* is seen when α-Cat is KD in the absence or presence of p35. In the absence of p35, the PC degenerates in this experiment, showing a genetic interaction between *α-CatRNAi(1)* and *puc-LacZ*. Scale bars, 100 μm. (**B**) Schematic illustrating the relationship between Puc and JNK. The phosphatase Puc dephosphorylates and therefore deactivates the JNK kinase. The presence of the puc mutant allele puc-lacZ is predicted to hyper-activate JNK in conjunction with a KD of α-Cat, which causes enhanced cell death as observed in (A). (**C**) Elevated expression of the Yki transcriptional reported *ex-lacZ* is observed upon KD of α-Cat in the absence or presence of p35. Tissue overgrowth is suppressed through the addition of a null mutant copy of *yki (yki^B5^)*. Scale bars, 100 μm. (**D**) Quantification of wing disc area in flies of indicated genotypes. Two-tailed, unpaired t-test used to determine statistical significance. ns (P>0.05), ****(P<0.0001), ***(P=0.0003).

Given previous findings in mammalian cells, we anticipated that the activation of Yki is responsible for the cell-autonomous overgrowth seen in *α-Cat-RNAi(1)* discs. Indeed, we found that the Yki transcriptional reporter *expanded (ex)-lacZ* is upregulated in the PC of *en>α-Cat-RNAi(1)* discs in addition to the previously reported [Yang et al., 2015], non-cell-autonomous upregulation in the AC (Figure 2C). Moreover, combined partial reduction of α-Cat and Yki (*α-Cat-RNAi(1), yki/+*) suppressed the *α-Cat-RNAi(1)* elicited overgrowth (Figure 2C,D). These findings indicate that a reduction of α-Cat causes a cell-autonomous upregulation of Yki activity. Interestingly, the *α-Cat-RNAi(1) yki/+* discs did not merely show suppressed overgrowth but were smaller than normal control discs with significantly reduced PCs. One important target of Yki is the cell death inhibitor Diap1 [Huang et al., 2005]. We speculate that under low Yki conditions and therefore reduced Diap1 expression, *en>α-Cat-RNAi(1)* discs have sufficient JNK activation to cause a substantial degeneration of the PC. Collectively, our findings suggest that the response to the loss of α-Cat function is governed by the differential activation of Yki and JNK signaling with Yki activation being more sensitive, causing tissue overgrowth under moderate α-Cat KD conditions.

### α-Catenin overexpression causes Yki-mediated tissue overgrowth

Our loss-of-function analysis suggested that the α-Cat expression level is important for the regulation of tissue growth. We therefore tested whether α-Cat overexpression has an impact on growth, expecting that overexpression would reduce growth given that lower than normal α-Cat levels enhance growth. Surprisingly, we found that α-Cat overexpression elicits overgrowth. We first noticed tissue overgrowth in the adult wing when α-Cat was overexpressed with *omb-Gal4* (Figure 3A,B). *omb-Gal4* is active in the central region of the developing wing pouch and hinge (Figure 1A). In contrast to the enlarged wings in *omb>α-Cat* flies, *omb>α-Cat-RNAi(1)* flies showed small wings with severe structural defects including notches and abnormal vein patterns, possibly the result of tissue degeneration at earlier stages of development (Figure 3A). Removing one copy of *yki* from *omb>α-Cat* flies normalized wing size (Figure 3A,B), suggesting that also overexpression of α-Cat enhances Yki activity.

**Figure 3:**
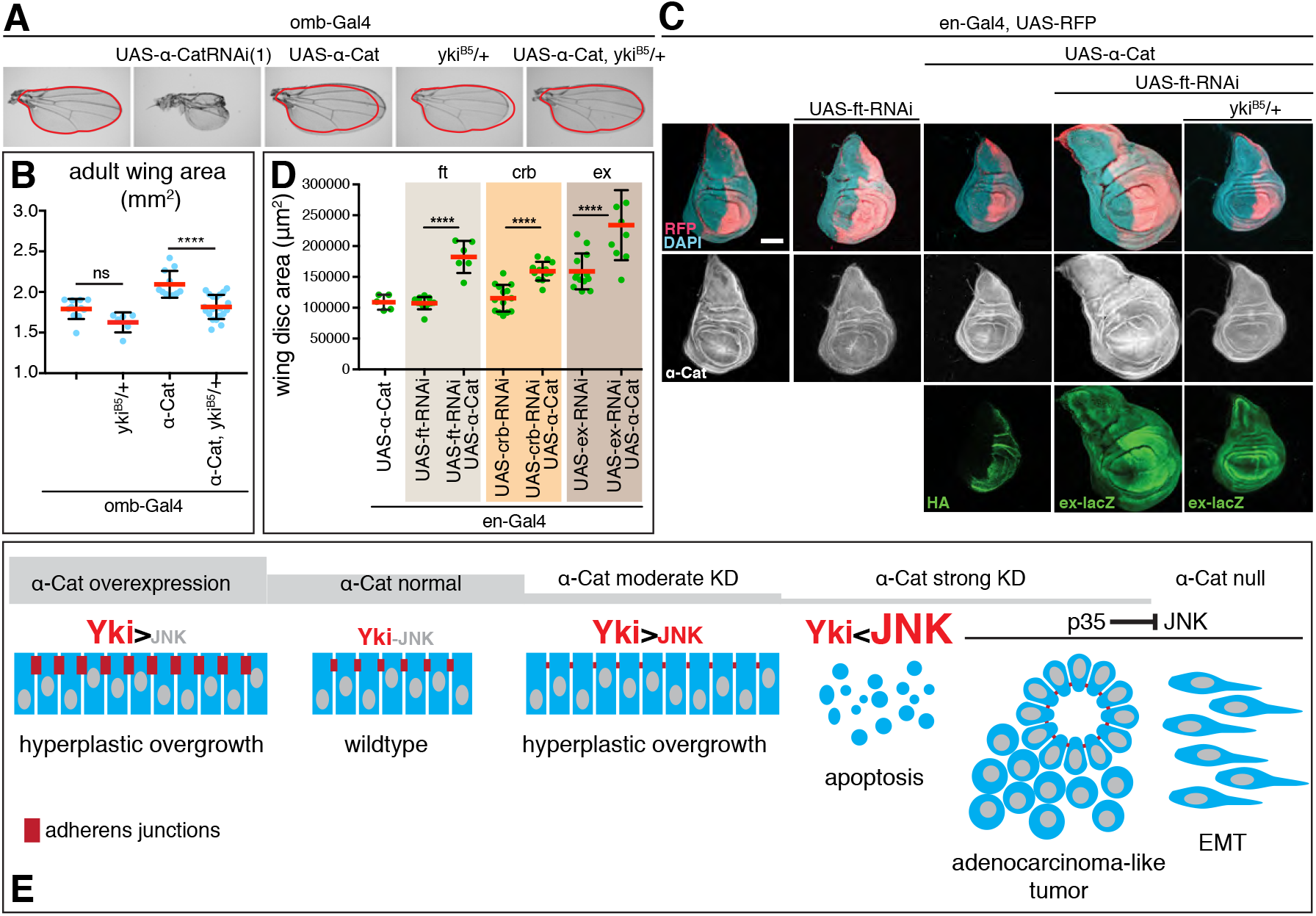
α-Cat overexpression activates Yki. (**A, B**) Overexpression of α-Cat causes an enlargement of the adult wing, which can be suppressed by one mutant copy of *yki*. Two-tailed, unpaired t-test used to determine statistical significance. ns(P>0.05), ****(P<0.0001). (**C, D**) Overexpression of α-Cat does not cause a significant enlargement of the late 3^rd^ larval instar wing disc. However, overexpression of α-Cat in conjunction with a KD of an upstream regulator of the Hippo/Yki pathway (Fat, shown in (C), Crb and Ex shown in Suppl. Figure S1; all three interactions quantified in (D)) show synergistic overgrowth phenotypes. Scale bars, 100 μm. Two-tailed, unpaired t-test used to determine statistical significance. ****(P<0.0001). (**E**) Illustration of the α-Cat phenotypic series and model for the differential activation of JNK and Yki signaling in response to changes in α-Cat levels.

We next examined larval wings discs overexpressing α-Cat with *en-Gal4* but did not detect any overgrowth (Figure 3C), suggesting that the enlarged adult wings result from overproliferation during pupal stages. We reasoned that overexpression of α-Cat alone may be insufficient to elicit excessive growth in a wild-type genetic background in larval wing discs. Hence, we sensitized the background by depletion of either Fat (Ft), Crb, or Ex, three positive upstream regulators of the Hippo pathway [Fulford et al., 2018]. The shRNAs used to KD Ft, Crb, and Ex caused little or no overgrowth on their own. However, when combined with α-Cat overexpression a synergistic overgrowth phenotype was observed in each case (Figure 3C,D; Suppl. Figure S1). For *ft-RNAi* we found that the interaction was suppressed by reducing Yki activity (Figure 3C). We conclude that α-Cat overexpression deregulates the Hippo/Yki pathway to cause overgrowth.

Taken together (Figure 3E), analysis of an α-Cat phenotypic series revealed that disruption of α-Cat can elicit Yki-dependent tissue overgrowth when α-Cat is either overexpressed or moderately down-regulated. When α-Cat levels further decline an increase in JNK signaling will overrite Yki-dependent overgrowth and cause tissue degeneration. In conjunction with a block of apoptosis, reduced levels of α-Cat caused an enhanced overgrowth phenotype, or, as α-Cat levels decline, disrupted epithelial polarity leading to an adenocarcinoma-like tumor, or, eventually, to a mesenchymal phenotype when α-Cat is lost completely.

### α-Catenin acts as an adherens junction protein to limit growth

Several mammalian studies have suggested the possibility that αE-catenin has cadherin-independent functions, possibly also in the regulation of growth [reviewed in Takeichi, 2018]. To address this question in Drosophila, we asked whether other core components of the CCC, DE-cadherin (DEcad) and p120catenin (p120), regulate tissue growth. We did not consider Arm as it is also a central player in the Wingless signaling pathway that is required for wing growth and development [Swarup and Verheyen, 2012]. Cell clones that lack DEcad did not develop in the wing disc similar to *α-Cat* null clones. To reproduce the effect of a moderate α-Cat KD, we partially depleted DEcad with RNAi in the presence of p35. *en>DEcad-RNAi p35* flies showed enlarged wing discs with cell-autonomous activation of Yki and JNK signaling similar to *en>α-Cat-RNAi(1)* or *en>α-Cat-RNAi(1) p35* (Figure 4A-C). Mature wings were not examined as *en>DEcad-RNAi* or *omb>DEcad-RNAi* flies did not survive to adulthood. Overexpression of DEcad *(omb>DEcad)* did result in slightly larger wings than controls similar to the overexpression of α-Cat (Figure 5B). We also examined *p120* null mutants, which show reduced DEcad levels at AJs but are not homozygous lethal [Myster et al., 2003] and found that both female (Figure 4D) and male (not shown) flies showed an increase in wing size when compared to controls. Thus, reduction or loss of α-Cat, DEcad, and p120 similarly affected wing size, suggesting that the level of the CCC plays an important role in growth regulation.

**Figure 4:**
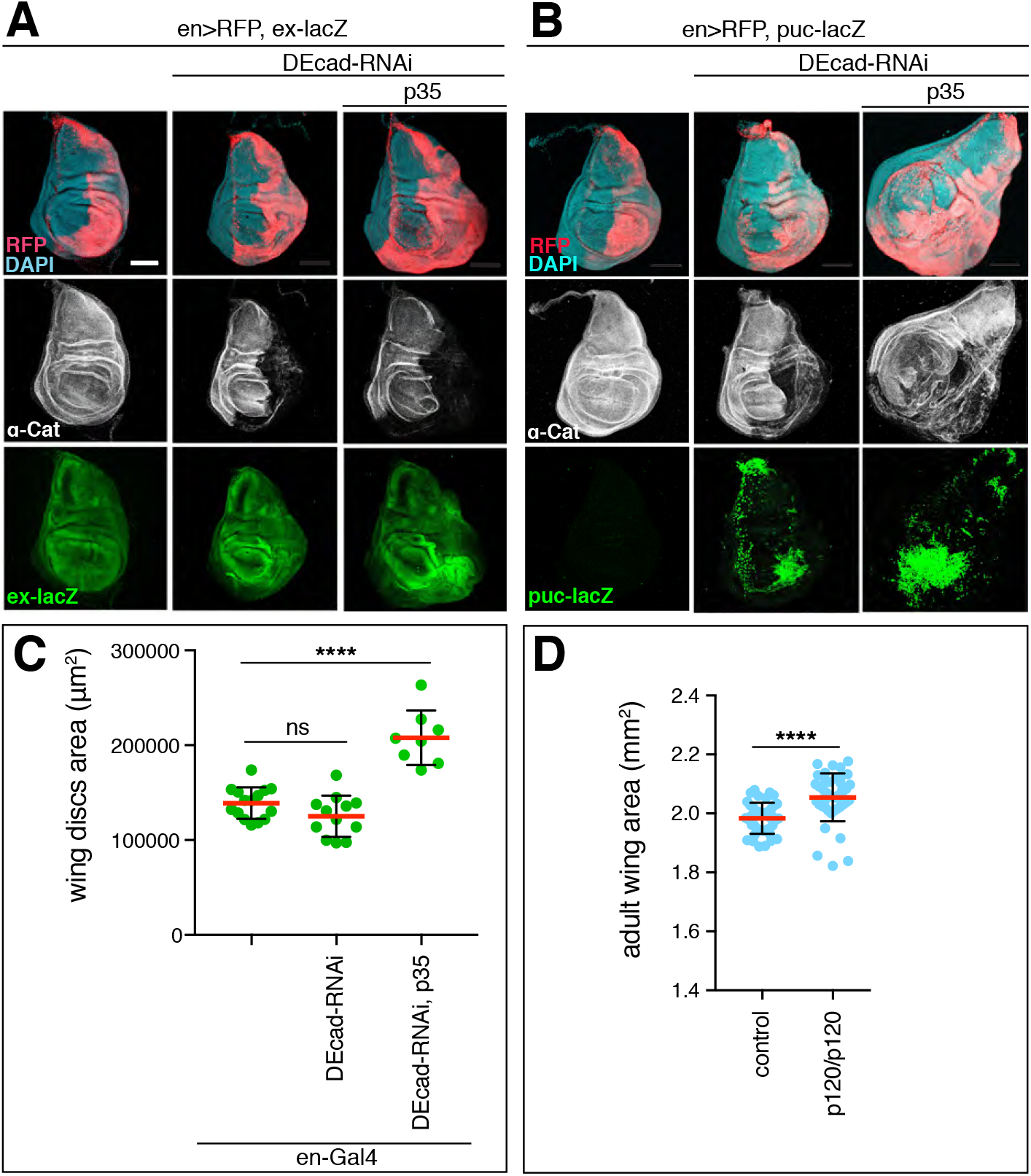
Reduction of DEcad and loss of p120catenin causes tissue overgrowth. (**A**) KD of DEcad enhances activation of the Yki transcriptional reporter *ex-LacZ*. KD of DEcad in conjunction with expression of p35 causes tissue overgrowth. Scale bars, 100 μm. (**B**) KD of DEcad with or without co-expression of p35 causes activation of the JNK transcriptional reporter *puc-LacZ*. KD of DEcad in conjunction with expression of p35 causes tissue overgrowth. Scale bars, 100 μm. (**C**) Quantification of late 3^rd^ larval instar wing disc area in DEcad KD flies. (**D**) Adult wing size of p120catenin null mutant flies compared to wildtype. Two-tailed, unpaired t-test used to determine statistical significance. ns (P>0.05), ****(P<0.0001).

**Figure 5:**
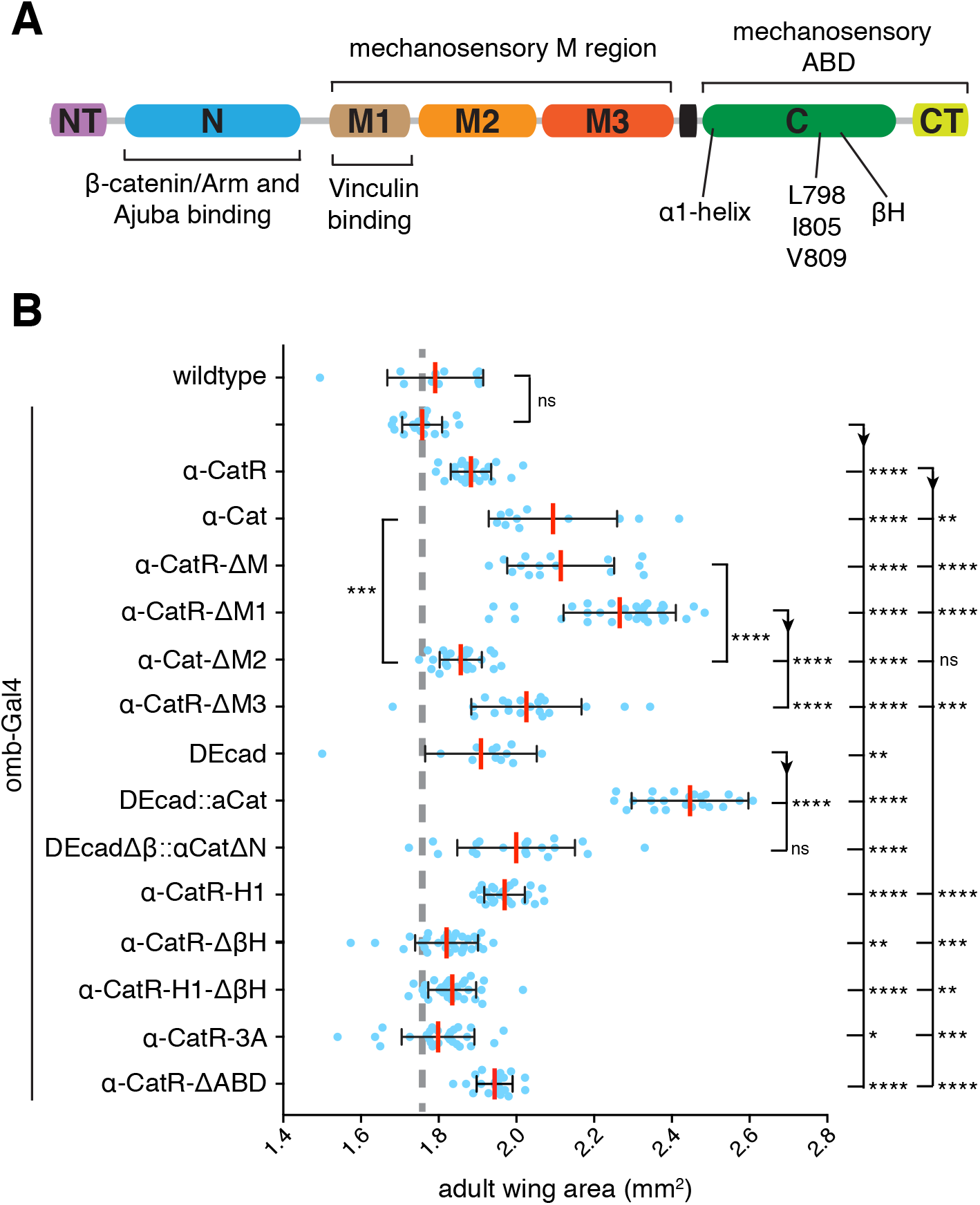
Overexpression of α-Cat mutant isoforms reveals domains important for limiting tissue growth. (**A**) α-Cat protein structure. (**B**) Adult wing area of flies overexpressing the indicated constructs with *omb-Gal4*. Two-tailed, unpaired t-test used to determine statistical significance. ns (P>0.05), *(P<0.05 and >0.01), **(P<0.01 and >0.001), ***(P<0.001 and >0.0001), ****(P<0.0001).

### The α-Cat N domain recruits Ajuba to AJs

α-catenin proteins are composed of a series of domains (Figure 5A). Loss of the the N domain which binds to β-catenin to couple α-catenin to the CCC and can dimerize, renders α-Cat nonfunctional because it is not recruited to the cadherin-β-catenin complex [Aberle et al., 1994; Pokutta and Weis, 2000; Desai et al., 2013]. However, a fusion protein in which α-Cat lacking the N domain is fused to DEcadΔβ (DEcadΔβ::αCatΔNT-N) can functionally replace the endogenous complex (Suppl. Figure 4) and allowed us to test for the requirement of N in growth regulation. As a control, we overexpressed DEcad:: αCat (a fusion of both full-length proteins) and observed a large increase in wing size (Figure 5B). This overgrowth was suppressed by reducing the activity of Yki (Suppl. Figure S2) suggesting that DEcad:: αCat acts by interfering with Hippo signaling. In contrast, DEcadΔβ::αCatΔNT-N overexpression was much less effective in inducing overgrowth compared to DEcad:: αCat and only showed an increase in wing size comparable to DEcad overexpression (Figure 5B). These results suggest that the NT-N domain plays an important role in growth regulation independent of its β-catenin-binding or dimerization activity.

Mammalian Ajuba can bind to the N-domain of αE-catenin [Marie et al., 2003]. Drosophila Jub and α-Cat can also form a complex, an interaction that was hypothesized to regulate Hippo/Yki signaling [Rauskolb et al., 2014]. As we found that Jub KD suppresses the overgrowth caused by DEcad::αCat overexpression (Suppl. Figure S2), we were curious to determine the level of junctional Jub in cells in which endogenous α-Cat was replaced by DEcadΔβ::αCatΔNT-N or DEcad::αCat. Monitoring Jub levels with a Jub::GFP construct expressed under control of its endogenous promoter [Sabino et al., 2011], we found that Jub levels in the AC and PC of control discs are similar as expected, whereas PC levels of Jub::GFP were reduced to 60% in *en>α-Cat-RNAi(1)* KD discs (Figure 6A,B,F). *en>α-Cat-RNAi(1)* KD cells expressing DEcad::αCat restored normal Jub levels to AJs (Figure 6C,F). In contrast, junctional Jub levels remained at 60% in *en>α-Cat-RNAi(1)* KD cells expressing DEcadΔβ::αCatΔNT-N similar to *en>α-Cat-RNAi(1)* cells (Figure 6,D,F). Examination of wing disc areas showed that expression of DEcad::αCat and DEcadΔβ::αCatΔNT-N in *en>α-Cat-RNAi(1)* KD discs were similar in size (Figure 6E). These findings indicate that NT-N domain is essential for Jub recruitment to AJs but also that the recruitment of Jub to AJs does not strongly correlate with the regulation of tissue growth in the larval disc expressing DEcad α-Cat fusion proteins.

**Figure 6:**
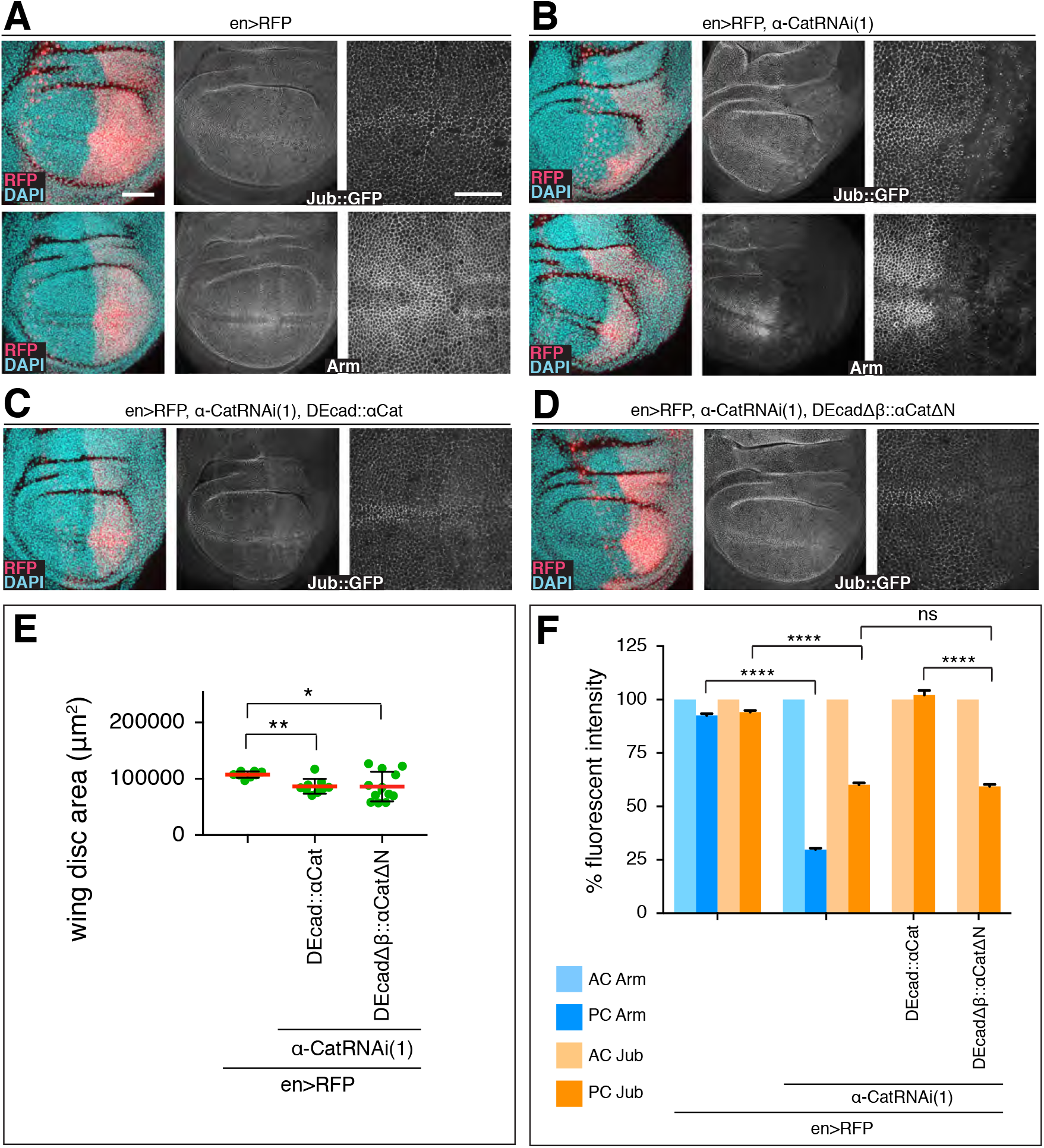
The α-Cat N domain recruits Jub to AJs. (**A**) Control late 3^rd^ larval instar discs expressing *en-Gal4, UAS-RFP* and either Jub::GFP controlled by its endogenous promoter (upper panels) or stained for Arm (lower panels). Nuclei are labeled with DAPI. Close-up images to the right show wing pouch area on both sides of the anterior-posterior compartment boundary. Scale bars, 50 μm and 25 μm. (**B**) Same markers as in (A) with *UAS-α-CatRNAi(1)* expression in posterior compartment. (**C**) Late 3^rd^ larval instar discs expressing Jub::GFP, *en-Gal4, UAS-RFP, UAS-α-CatRNAi(1)*, and *UAS-DEcad::αCat*. (**D**) Late 3^rd^ larval instar discs expressing Jub::GFP, *en-Gal4, UAS-RFP, UAS-α-CatRNAi(1)*, and *UAS-DEcadΔβ::αCatΔN*. (**E**) Quantification of wing disc area of flies of indicated genotypes. Two-tailed, unpaired t-test used to determine statistical significance; ****(P<0.0001). (**F**) Comparison of relative fluorescent intensities between anterior compartment (AC) and posterior compartment (PC) for Jub::GFP and Arm. AC values were normalized to 100%. N=200-500 cells from three wing discs. Mann Whitney test was used to determine statistical significance; ****(P<0.0001), ns (P>0.05).

### The M region of α-Cat modulates tissue growth

We took advantage of the Yki-dependent increase in adult wing size upon α-Cat overexpression to assess the function of different α-Cat domains in tissue growth. For these tests we used constructs we published previously [Sarpal et al., 2012; Desai et al., 2013; Ishiyama et al., 2018] and a new set of constructs that were rendered resistant to *α-Cat-RNAi(1)* and *α-Cat-RNAi(2)* and will be designated as α-CatR-XX. Both α-Cat and α-CatR encode wildtype proteins. Although all constructs were expressed from the same genomic insertion site under UAS control we noted that α-CatR-based constructs were expressed at a lower level than α-Cat-based constructs (Suppl. Figure S3). When expressed in *en>α-Cat-RNAi(2)* PC cells α-CatR levels were ~2/3 of normal (Suppl. Figure S3). α-Cat and α-CatR rescued *α-Cat* mutants to adulthood [Desai et al., 2013; Suppl. Figure S4], and α-CatR fully rescued integrity and growth of *α-Cat-RNAi(1)* or *α-Cat-RNAi(2)* expressing wing epithelium (see below). For statistical analyses, we have compared α-CatR mutant constructs to α-CatR and all other mutant constructs to α-Cat. Consistent with the lower expression level, we found that the increase in wing size is smaller with *omb>α-CatR* than *omb>α-Cat* (Figure 5B).

Mechanical tension can regulate tissue growth in the wing disc such that high tension elicits more growth than low tension [Eder et al., 2017; Diaz de la Loza and Thompson, 2017]. α-catenin is thought to respond to actomyosin generated tension through two conformational changes in the M region and the ABD, respectively. Tension reveals cryptic binding site for the actin-binding protein Vinculin and perhaps other partners in the M region [Yonemura et al., 2010; Mege and Ishiyama, 2017; Ishiyama et al., 2018]. The M region-based mechanosensing of α-Cat was hypothesized to mediate the cytoskeletal tension-dependent recruitment of Jub to AJs to negatively regulate tissue growth [Rauskolb et al., 2014]. To test this idea we deleted the entire M region from α-CatR (α-CatR-ΔM; Suppl Figure S4) and investigated *en>α-Cat-RNAi(2) α-CatR-ΔM* flies. The results were striking as α-CatR-ΔM was not only recruited to AJs in the wing epithelium (Suppl. Figure S5) but also fully restored a normal disc in both epithelial organization and size similar to α-CatR (Figure 7A,C). Moreover, flies survived to adulthood with normal wings (Figure 7E). These data show that the M region, and with it M region-based mechanosensing is not essential for growth regulation in the wing epithelium.

**Figure 7:**
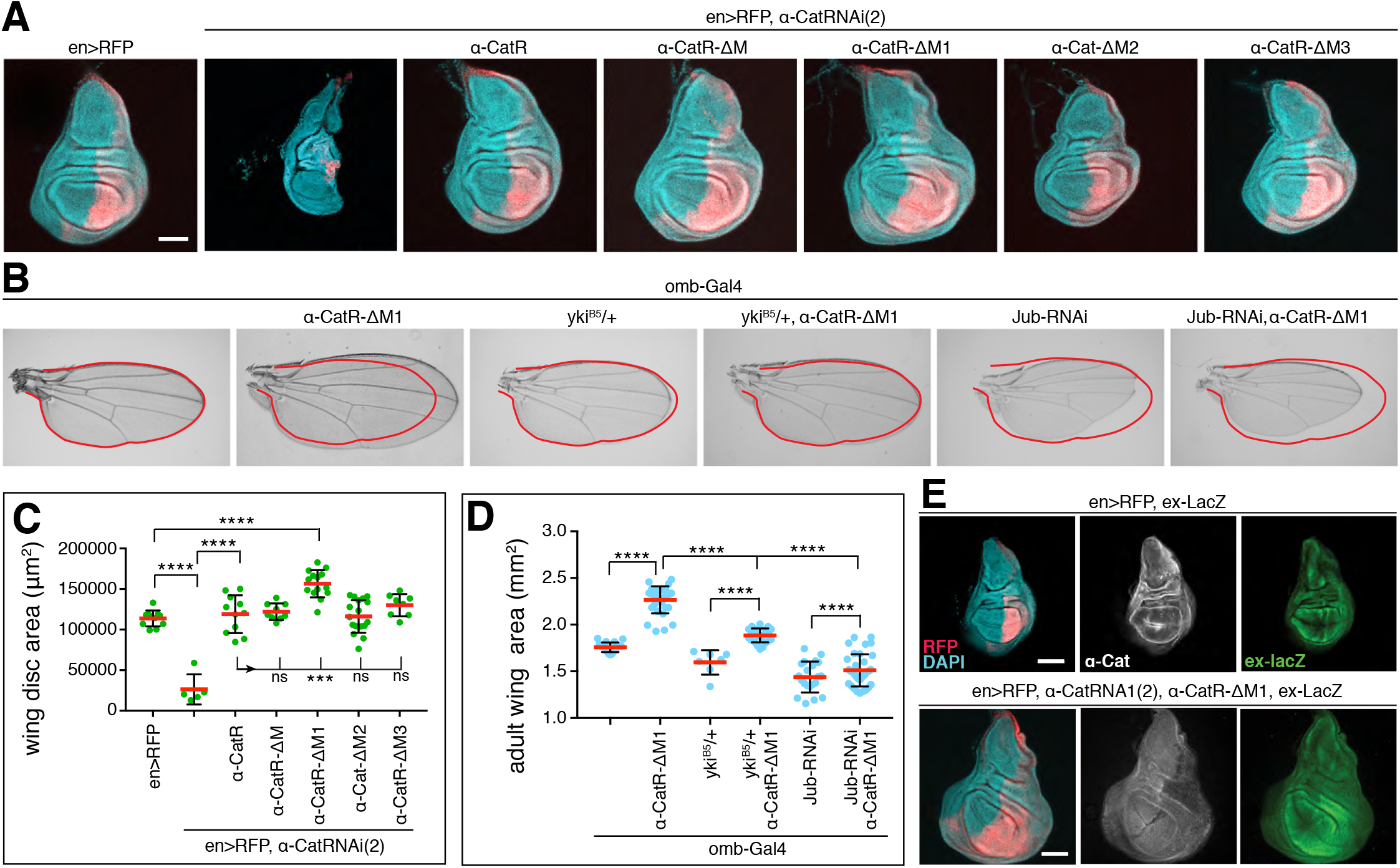
The M1 domain of α-Cat limits Jub- and Yki-dependent tissue overgrowth. (**A**) Late 3^rd^ larval instar wing discs of indicated genotypes labeled with DAPI and posterior compartment marked by RFP. α-Cat was depleted in PC with *α-Cat-RNAi(2)*. Scale bars, 100 μm. (**B**) Adult wings of flies of indicated genotypes. Both *yki^B5^/+* and Jub-RNAi suppress the overgrowth elicited by expression of α-CatR-ΔM1. (**C**) Quantification of late 3^rd^ larval instar wing disc area corresponding to (A) of flies of indicated genotypes. Two-tailed, unpaired t-test used to determine statistical significance; ****(P<0.0001). (**D**) Quantification of adult wing area corresponding to (B) of flies of indicated genotypes. Twotailed, unpaired t-test used to determine statistical significance; ****(P<0.0001). (**E**) Wing of control (*en>RFP*) and *en>RFP, α-Cat-RNAi(2), α-CatR-ΔM* adult fly. (**F**) Late 3^rd^ larval instar wing disc of the indicated genotype showing the failure of *α-CatR-ΔM1* to rescue tissue overgrowth in an α-Cat compromised background (compare to Figure 2C).

*omb>α-CatR-ΔM* adult flies had wings that were overgrown similar to *omb>α-CatR* flies (Figure 5B), in line with the notion that α-CatR-ΔM functions like α-Cat in wing development. We were therefore surprised to find that α-Cat isoforms in which individual domains of the M region were deleted (which are all enriched at AJs; Figure 5B and Suppl. Figure S5), showed distinctly different responses. The M region is composed of three α-helical bundles, M1, M2, and M3. *omb>α-CatR-ΔM1* flies had wings that were larger than *omb-Gal4/+, omb>α-CatR* or *omb>α-Cat* controls and larger than wings of *omb>α-Cat-ΔM* flies. In contrast, *omb>α-Cat-ΔM2* wings were smaller than *omb>α-Cat* or *omb>α-Cat-ΔM* and only slightly larger than *omb-Gal4/+* controls. *omb>α-CatR-ΔM3* flies had wings smaller than those of *omb>α-CatR-ΔM1* flies and only slightly larger than *omb>α-CatR* control flies. We note again that we compared shRNA resistant “R” constructs against a *α-CatR* control and non-resistant constructs against *α-Cat*. The data shown in Figure 5B therefore likely underestimate the difference between *α-CatR-ΔM1* and *α-Cat-ΔM2* as R constructs have a lower expression level compared to non-R. These findings suggest that M1 and to a lesser extent M3 normally act to limit the ability of α-Cat to stimulate growth, whereas M2 is important for overexpressed α-Cat to stimulate growth. The opposing effects of deleting M1 and M3 versus M2 may explain why α-CatR-ΔM behaves similarly to full-length controls.

*en>α-Cat-RNAi(2) α-CatR-ΔM* imaginal disc were normal as mentioned above. In contrast, *en>α-Cat-RNAi(2) α-CatR-ΔM1* showed hyperplastic overgrowth similar to discs with a moderate KD of α-Cat (Figure 7A,C), whereas *en>α-Cat-RNAi(2) α-Cat-ΔM2* and *en>α-Cat-RNAi(2) α-CatR-ΔM3* discs were normal in size (Figure 7A,C). Thus, all tested deletion constructs of the M region rescue epithelial polarity and growth defects of strong α-Cat KD conditions in the wing epithelium, with the exception of α-CatR-ΔM1 which rescued only epithelial polarity but not hyperplastic overgrowth. Consistently, we observed enhanced activity of the Yki reporter *ex-LacZ* in *en>α-Cat-RNAi(2) α-CatR-ΔM1* discs (Figure 7F), and we found that α-CatR-ΔM1 elicited overgrowth is suppressed by reducing *yki* (Figure 7B,D). These data together with the findings from overexpression experiments suggest that M1 has an important role in limiting tissue growth.

### The M1 domain of α-Cat restricts the N-domain-dependent recruitment of Ajuba

We also noticed that the overgrowth caused by α-Cat-ΔM1 expression is suppressed by depletion of Jub (Figure 7B,D). We asked therefore whether Jub localization is affected in cells expressing M region deletion constructs. We quantified the junctional concentration of Jub::GFP in AC control cells versus PC cells expressing *en> α-Cat-RNAi(2)* and an α-Cat construct. We also determined the levels of the CCC by quantifying junctional Arm. The latter was done to find out whether potential changes in Jub levels reflect changes in CCC concentration given that Jub is a direct binding partner of α-Cat [Marie et al., 2003; Rauskolb et al., 2014], and that this interaction is crucial for junctional localization of Jub (Figure 6). Alternatively, a lack of correlation in changes of junctional Jub and Arm would indicate that the tested α-Cat mutation affects the Jub-α-Cat interaction more directly.

In *en> α-Cat-RNAi(2) α-CatR* control cells we found that levels of Arm were reduced to 66%, consistent with the reduced protein levels of α-CatR compared to endogenous α-Cat (Suppl. Figure S3). Surprisingly, Jub::GFP was not reduced correspondingly but increased to 119% compared to AC cells (Figure 8A,F), indicating that junctional Jub does not directly follow CCC level, but that other mechanisms and possibly other binding partners could contribute to junctional localization of Jub. Similar to α-CatR, M region deletions caused lower Arm levels as seen in control cells. α-CatR-ΔM expressing cells showed only 32% of normal Arm levels but, interestingly, also a ~30% increase in junctional Jub::GFP (131%; Figure 8B,F). α-CatR-ΔM3 had Arm levels of 64% similar to α-CatR but Jub was not enhanced (98%; Figure 8E,F). α-Cat-ΔM2 had the lowest Arm levels with 22% and also showed a reduction of Jub::GFP to 78% (Figure 8D,F). This is consistent with our previous findings that the M2 region has a strong impact on junctional stability [Desai et al., 2013]. The most striking finding was the more than 4-fold increase (468% of normal) in junctional Jub::GFP levels observed with α-CatR-ΔM1 with Arm levels at only 59% (Figure 8C,F).

**Figure 8:**
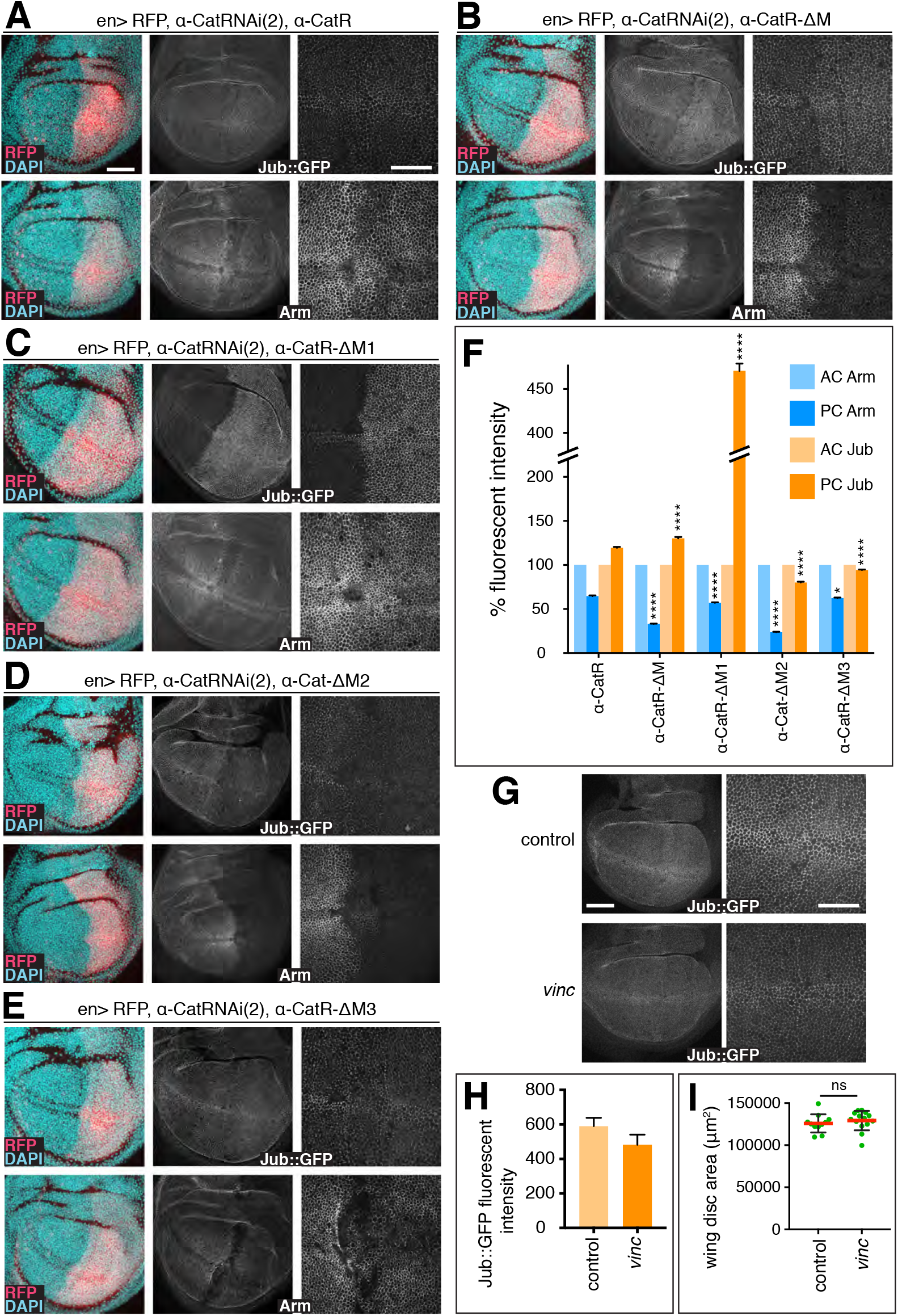
The M1 domain of α-Cat limits recruitment of Jub to AJs. (**A-E**) Late 3^rd^ larval instar discs expressing *en-Gal4, UAS-RFP, UAS-α-CatRNAi(2)* and either *UAS-α-CatR* (A), *UAS-α-Cat-ΔM*(B), *UAS-α-CatR-ΔM1* (C), *UAS-α-Cat-ΔM2* (D), or *UAS-α-CatR-ΔM3* (E). Disc express either Jub::GFP controlled by its endogenous promoter (upper panels) or are stained for Arm (lower panels). Nuclei are labeled with DAPI. Close-up images to the right show wing pouch area on both sides of the anterior-posterior compartment boundary. Scale bars, 50 μm and 25 μm. (**F**) Comparison of relative fluorescent intensities between anterior compartment (AC) and posterior compartment (PC) for Jub::GFP (N=500-1000 cells from five wing discs) and Arm (N=500-1000 cells from four to six wing discs). AC values were normalized to 100%. Mann Whitney test was used to determine statistical significance. ****(P<0.0001), *(P=.0258) (**G**) Jub::GFP expression in control and *vinc* null mutant (*vinc102.1/vinc102.1*) late 3^rd^ larval wing discs. (**H**) *vinc* null mutant and control late 3^rd^ larval wing discs have the size. Two-tailed, unpaired t-test was used to determine statistical significance. ns (P>0.05) (**I**) Quantification of Jub::GFP fluorescent intensities (arbitrary units) in control and *vinc* null mutant discs (N>900 cells from five wing discs).

Together, these findings indicate that the M region, and all three M domains are required to stabilize the CCC. However, loss of the M region does not deplete the CCC sufficiently to compromise epithelial integrity in the wing disc. In each case, including the α-CatR control, we find that Jub levels do not correspond to the observed reduction in Arm levels and in case of α-CatR and α-CatR-ΔM moderately exceed wild-type levels. Moreover, the striking increase in Jub enrichment at AJs with α-CatR-ΔM1 indicates that M1 strongly limits the junctional recruitment or stability of Jub. This explains why α-CatR-ΔM1 fails to rescue the tissue overgrowth resulting form α-Cat depletion (Figure 7), elicits tissue overgrowth when overexpressed (Figure 5B), and likely implies that Jub recruitment to α-Cat has been decoupled from tissue tension. These conclusions are supported by the observed suppression of α-Cat-ΔM1 elicited overgrowth when Yki or Jub were reduced (Figure 7B,D). The fact that the loss of the entire M region does not adversely affect the regulation of tissue growth may be the result of compensatory activities of M1, which normally limits junctional localization of Jub, and M2 and M3, which normally appear to enhance junctional localization of Jub (Figure 8F). We conclude that the M region of α-Cat has complex activities that contribute to the regulation of AJ stability and the Jub recruitment-dependent regulation of tissue growth.

### Vinculin is not required for the α-CatR-ΔM1 elicited Jub hyper-recruitment or tissue overgrowth

The striking impact of deleting M1 on Jub junctional localization prompted us to ask whether Vinculin is involved as Vinculin is a well-established binding partner of M1 [Watabe-Uchida et al., 1988; Yonemura et al., 2010; Choi et al., 2012; Yao et al., 2014]. Drosophila *Vinculin (Vinc)* null mutants are homozygous viable and fertile [Alatortsev et al., 1997; Klapholz et al., 2015]. We found Jub::GFP levels were not increased in *Vinc* mutant 3rd larval wing discs compared to controls (Figure 8G,H), and wing discs were not enlarged compared to wild-type (Figure 8I). These findings suggest that Vinc does not play an essential role in tissue growth of the wing disc epithelium and that the mechanism explaining how M1 could limit Jub recruitment to AJs does not appear to involve its interaction with Vinc.

### Mechanosensing of the α-Cat actin-binding domain is required for normal growth regulation

We recently identified a molecular mechanism for the apparent catch-bond behavior of α-catenin ABD [Buckley et al, 2014; Ishiyama et al., 2018]. The first α-helix of the α-catenin ABD, α1-helix, undergoes a conformational change under physiological tension, consequently revealing an enhanced actin-binding interface and a small β-sheet hairpin loop (βH) that facilitates ABD dimerization (see Figure 5A for the location of sequence elements). In vitro actin-binding assays suggest that a mutation in the α1-helix that compromises its helical structure (H1) or deletion of the α1-helix enhance actin-binding and bundling activities, whereas deletion of βH (ΔβH) or a double mutant combination of H1 and ΔβH reduces actin-binding [Ishiyama et al., 2018]. Thus, in contrast to removing the ABD, which renders α-Cat non-functional [Desai et al., 2013; Ishiyama et al., 2018], H1 and ΔβH provide more subtle changes in ABD activity that either enhance or weaken the direct interaction of α-catenin with F-actin. The corresponding mutant fly isoforms (α-CatR-H1, α-CatR-ΔβH, α-CatR-H1-ΔβH), as well as isoforms that compromise the actin-binding interface (L798A+I805A+V809A = α-CatR-3A), or lack ABD (α-CatR-ΔABD) were all effectively recruited to AJs (Suppl. Figure S5) [Ishiyama et al., 2018]. α-CatR-3A, α-CatR-ΔABD, and α-CatR-ΔβH did not rescue the embryonic lethal phenotype of *α-Cat* zygotic mutants. α-CatR-H1 and α-CatR-H1-ΔβH showed some rescue activity, with α-CatR-H1-ΔβH having a better rescue than α-CatR-H1. This is consistent with the notion that the H1 mutation, which enhances actin-binding, and the ΔβH mutation, which reduces actin binding, may compensate for each other to some degree. However, most α-CatR-H1 and α-CatR-H1-ΔβH rescued embryos still died as embryos or during early larval stages in contrast to α-CatR expressing animals, which rescues *α-Cat* mutants to adults [Ishiyama et al., 2018].

Overexpression of ABD mutant α-Cat proteins in the wing showed relatively minor effects and adult wing size with expression of α-CatR-3A, α-CatR-ΔβH, α-CatR-H1-ΔβH slightly smaller wings than seen in α-CatR controls, and expression of α-CatR-ΔABD and α-CatR-H1 led to small increases in wing size (Figure 5B). We next expressed α-Cat ABD mutants together with *en>α-Cat-RNAi(2)* in the wing imaginal disc. Whereas α-CatR-H1, α-CatR-ΔβH, and α-CatR-H1-ΔβH had substantial rescue activities (Figure 9A,B) compared to *en> α-Cat-RNAi(2)* discs that had lost most of their PCs (Figure 1E and 7A), we could not recover 3^rd^ instar larvae with α-CatR-3A. We therefore tested α-CatR-3A as well as α-CatR-ΔABD in a background expressing the weaker *α-Cat-RNAi(1)* KD line. *en>α-Cat-RNAi(1)* on its own caused hyperplastic overgrowth (Figure 1C,E). In contrast, in conjunction with α-CatR-3A or α-CatR-ΔABD we recovered discs that essentially lacked a PC (Figure 9A,B) similar to discs expressing *en>α-Cat-RNAi(2)*. These observations further emphasize that direct actin binding is critical for α-Cat function in vivo [Desai et al., 2013; Ishiyama et al., 2018], and suggest that α-CatR-3A and α-CatR-ΔABD act as dominant-negative isoforms of α-Cat, interfering with the function of the endogenous protein that is not removed through KD by either *α-Cat-RNAi(1)* or *α-Cat-RNAi(2)*.

**Figure 9:**
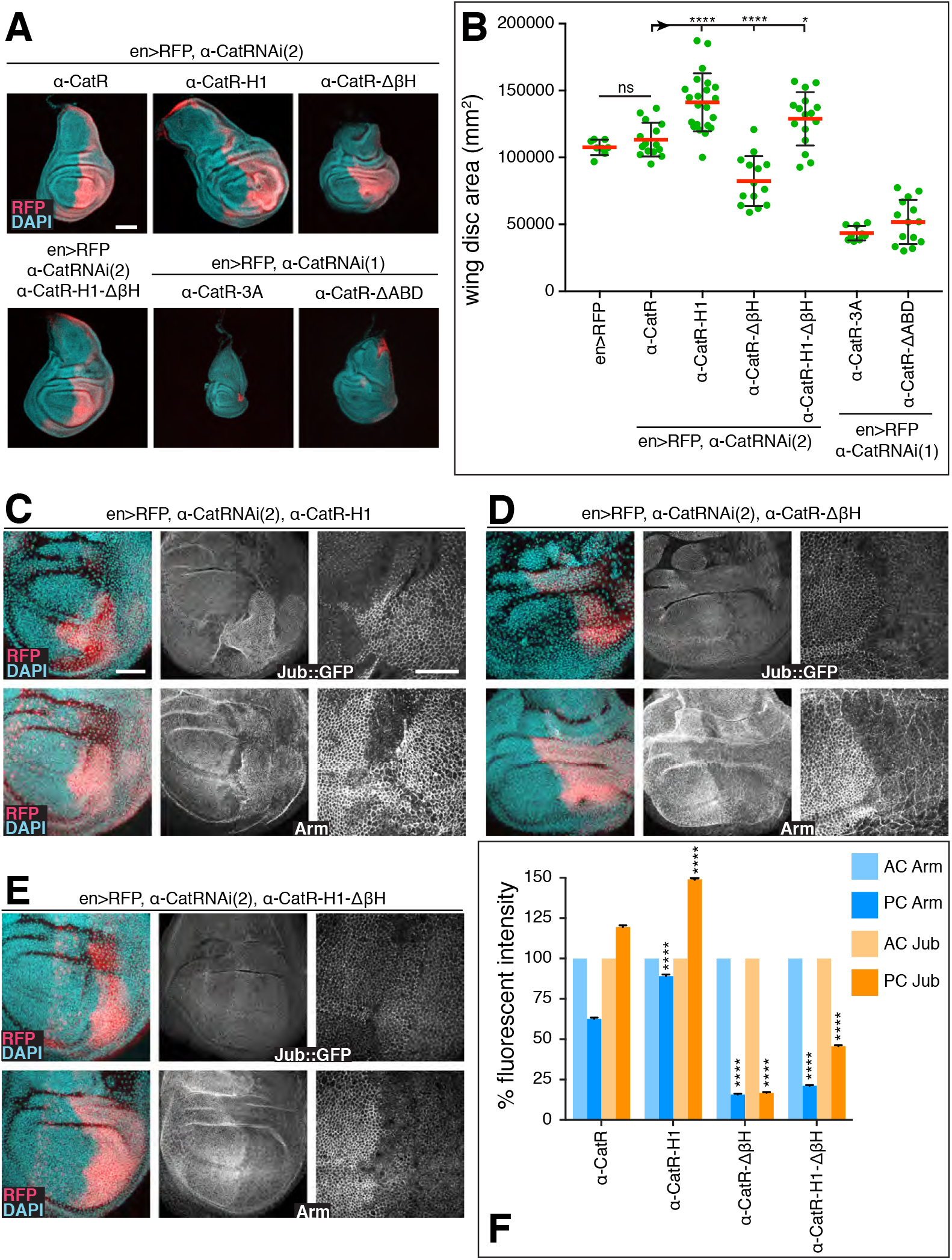
The mechanosensory α1-helix of the α-Cat ABD regulates tissue growth. (**A, B**) Sample discs (A) and quantification of disc area (B) of late 3^rd^ larval instar wing discs of the indicated genotypes. Note that UAS-α-CatR-H1 expression in *en-Gal4, UAS-α-CatRNAi(2)* discs causes hyperplastic overgrowth. α-CatR-3A and α-CatR-ΔABD expression enhanced the phenotype of *en>α-CatRNAi(1)* discs. 3^rd^ larval discs expressing α-CatR-3A in a *en>α-CatRNAi(2)* background did not develop. Scale bars, 100um. (**C-E**) Late 3^rd^ larval instar discs expressing *en-Gal4, UAS-RFP, UAS-α-CatRNAi(2)* and either *UAS-α-CatR-H1* (C), *UAS-α-Cat-ΔβH* (D), or *UAS-α-CatR-H1 -ΔβH* (E). Disc express either Jub::GFP controlled by its endogenous promoter (upper panels) or are stained for Arm (lower panels). Nuclei are labeled with DAPI. Close-up images to the right show wing pouch area on both sides of the anterior-posterior compartment boundary. Scale bars, 50 μm and 25 μm. (**F**) Comparison of relative fluorescent intensities between anterior compartment (AC) and posterior compartment (PC) for Jub::GFP (N=500-1000 cells from five wing discs??? check) and Arm (N=500 - 1000 cells from three to four wing discs). AC values were normalized to 100%. Mann Whitney test was used to determine statistical significance. ****(P<0.0001)

α-CatR-H1, α-CatR-ΔβH, and α-CatR-H1-ΔβH showed intriguing differences in their ability to rescue the depletion of the endogenous protein. *en>α-Cat-RNAi(2) α-CatR-ΔβH* discs showed normal epithelial organization of the PC but overall disc size remained smaller than of α-CatR controls (Figure 9A,B). *en>α-Cat-RNAi(2) α-CatR-H1* discs were overgrown similar to *en>α-Cat-RNAi(1)* discs suggesting that α-CatR-H1 supports normal epithelial polarity but fails in growth regulation. Finally, *en>α-Cat-RNAi(2) α-CatR-H1-ΔβH* discs showed an amount of tissue overgrowth intermediate between α-CatR-H1 and α-CatR (Figure 9A,B), suggesting that α-CatR-H1-ΔβH supports epithelial integrity and, to a larger extent than α-CatR-H1, the normal growth regulatory function of α-Cat.

In comparing junctional Jub::GFP and Arm levels in *en>α-Cat-RNAi(2)* discs expressing α-CatR-H1, α-CatR-ΔβH, or α-CatR-H1-ΔβH we found that Arm was reduced to 18% and 21% in PC compared to AC cells in α-CatR-ΔβH and α-CatR-H1ΔβH expressing discs, respectively (Figure 9D-F). This suggests that these mutant forms do not support normal stability of the CCC, and consequently cause significantly lower junctional Jub levels of 17% and 46% compared to controls. In contrast, α-CatR-H1 discs displayed higher than control levels of Arm (88% versus 63%) and Jub (149% versus 119%) (Figure 9C,F). This is interesting as it suggests that the enhanced actin-binding activity observed for α-CatR-H1 in vitro [Ishiyama et al., 2018] causes the formation of more stable AJs in vivo. The elevated recruitment of Jub to AJs likely accounts for the observed tissue overgrowth of *en>α-Cat-RNAi(2) α-CatR-H1* discs (Figure 9A,B). Together, these results suggest that mechanosensing of the α-Cat ABD and the precise mechanism of the α-Cat F-actin association are important components of the AJ-dependent regulation of tissue growth.

### Loss of α-Cat mechanosensing does not change tissue tension

The tension generated by cytoskeletal forces recruits Jub to AJs [Razzell et al 2018., Rauskolb et al., 2014] and modulates cell proliferation in the wing disc [Eder et al., 2016]. That some α-Cat mutant isoforms induce increased tension is a possible explanation for the excessive recruitment of Jub. We were wondering therefore whether mutations in α-Cat that eliminated M region or α1-helix-dependent mechanosensing led to overt changes in tissue tension that could impact growth. To address this question, we determined the initial recoil velocity after laser ablation of cell junctions in experimental PC cells versus control AC cells (Figure 10A). We tested α-CatR, α-CatR-ΔM, α-CatR-ΔM1, and α-CatR-H1 in an *en>α-Cat-RNAi(2)* Jub::GFP background. These tests did not reveal significant differences in tissue tension in comparing AC control cells to PC experimental cells (Figure 10B), suggesting that the impact the tested α-Cat mutant isoforms have on growth is not through a change in tissue tension. We noted, however, that wound healing was impaired in α-CatR-H1 expressing PC cells (Figure 10C) similar as previously observed in the embryonic epidermis [Ishiyama et al., 2018]. These findings support the view that α-Cat acts as a mechanosensor that translates cytoskeletal tension into biochemical signals that limit tissue growth. Moreover, in the case of α-CatR-ΔM1 rescued PC cells, Jub junctional recruitment is highly enriched (Figure 8C,F) even as tension tends to be lower than in control PC cells (Figure 10C). This suggests that the ability for Jub recruitment to respond to different tension states has been lost with the removal of the M1 domain, and instead Jub is constitutively recruited to AJs regardless of the state of tension.

**Figure 10:**
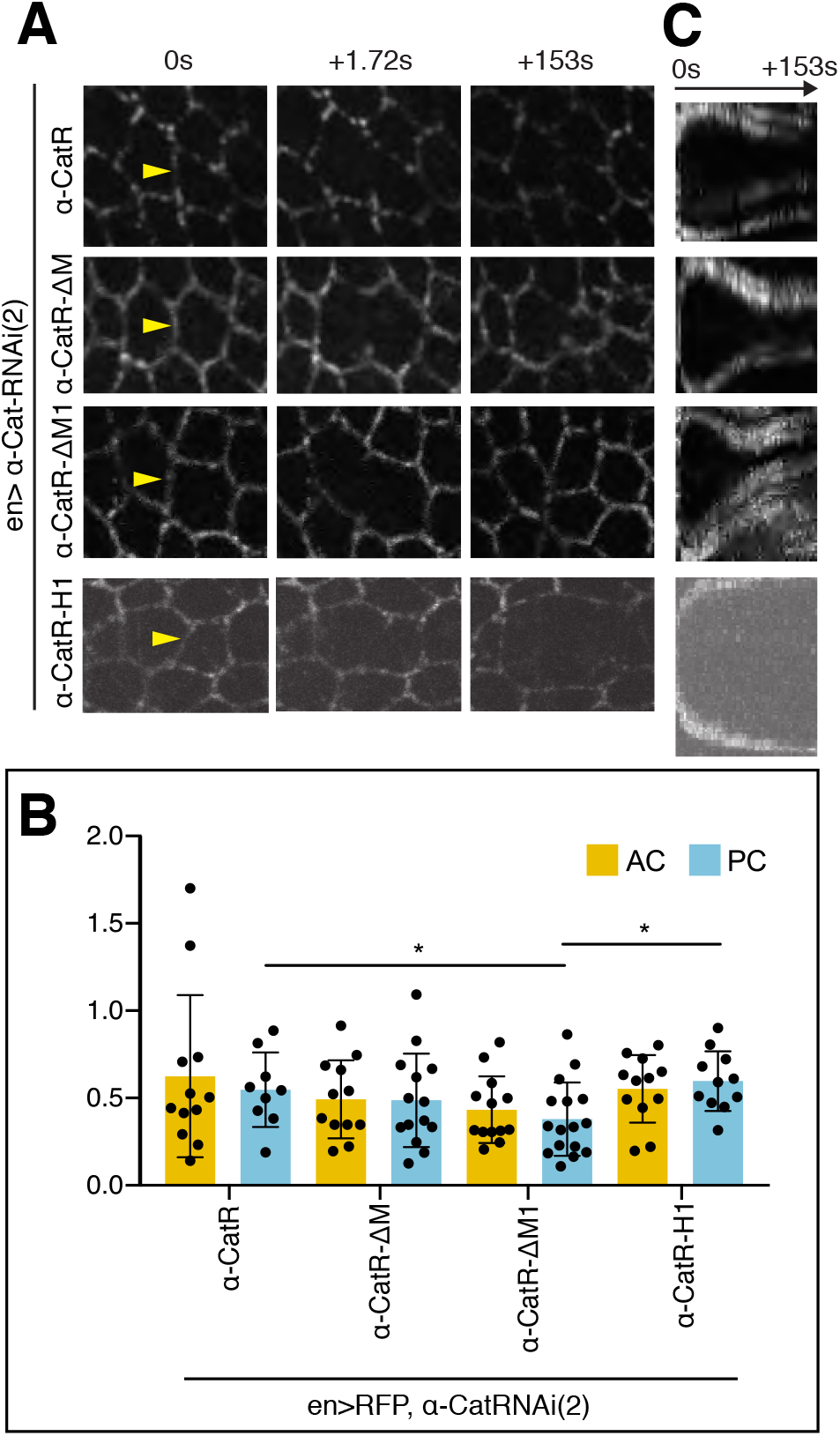
α-Cat mechanosensory mutants do not alter tissue tension. (**A**) Late 3^rd^ larval instar wing imaginal discs expressing Jub::GFP, *en-Gal4, UAS-RFP, UAS-α-CatRNAi(2)*, and either *UAS-α-CatR, UAS-α-Cat-ΔM, UAS-α-CatR-ΔM1*, or *UAS-α-CatR-H1*. Junctions were cut with an intense laser pulse (see Materials and Methods) at the sites indicated by the arrowheads. Recoil velocity (B) and wound closure (B) were determined from movie sequences. (**B**) Quantification of initial recoil velocities of cell edges after laser ablation in wing discs with indicated genotypes. Values of posterior compartments (PC) are compared with anterior compartment (AC) control cell values with 3-5 discs used per genotype. No significant differences between anterior and posterior cells were observed. Two-tailed, unpaired t-test was used to determine statistical significance; *(≤ 0.0332). (**C**) Kymographs recording wound closure after laser cut. α-CatR, α-Cat-ΔM, and CatR-ΔM1 support normal wound closure dynamics whereas α-CatR-H1 expressing discs cells display delays in wound healing.

## Discussion

One key property of α-Cat that determines its impact on tissue growth, leading either to overgrowth or tissue degeneration, is the level of α-Cat present at AJs. Through an analysis of an α-Cat phenotypic series we could document a role for Drosophila α-Cat as a negative regulator of tissue growth. Moderate α-Cat overexpression or moderate depletion of α-Cat or DEcad causes Yki activation and overproliferation. Moreover, consistent with our present results, previous analysis of strong loss-of-function conditions for α-Cat, DEcad, or Arm led to the conclusion that a loss of the CCC reduces Yki activity and causes JNK-meditated tissue degeneration [Yang et al., 2015]. This was at odds with some mammalian studies where the loss of E-cadherin or αE-catenin causes YAP or TAZ-dependent overgrowth in a number of different tissues and cell lines [for review see Gumbiner and Kim, 2014; Karaman and Halder et al., 2018]. Our data showing that α-Cat and DEcad limit Yki activity and tissue growth in Drosophila similar to mammalian tissues suggest a conserved functional relationship between AJs and the regulation of Yki/YAP/TAZ activity. Only when the Drosophila CCC is strongly depleted does a substantive activation of JNK signaling override Yki activation, causing tissue degeneration. This could potentially involve a direct inhibition of Yki by JNK [Enomoto et al., 2015].

*p120* null mutants, which are homozygous viable [Mysters et al., 2003], also have enlarged adult wings [see also Iyer et al., 2019] consistent with an observed moderate reduction in CCC levels. p120, E-cadherin, and αE-catenin were recently identified as prominent haploinsufficient tumor suppressors in mammalian models for intestinal cancer [Short et al., 2017]. Notably, complete loss of mammalian p120, which has a more dramatic impact on membrane abundance of the CCC than the loss of its Drosophila homolog, causes a disruption of the intestinal epithelium and cell death. Therefore, both the Drosophila wing epithelium and the mammalian intestinal epithelium respond to a moderate loss of the CCC with overgrowth whereas a strong reduction causes cell death. Hippo/YAP signaling has been implicated in tumor formation in the mammalian intestine [Yu et al., 2015]. However, whether it acts downstream of the CCC in this tissue remains unknown. The cell death observed upon the complete loss of p120 in intestinal cells that also lack the tumor suppressor APC is effectively blocked by inhibition of Rho kinase, suggesting that Rho1 signaling is required for apoptosis [Short et al., 2017]. In Drosophila, the RhoGEF2-Rho1-Rho kinase-Myosin II pathway was linked to JNK activation [Neisch et al., 2010; Khoo et al., 2013], raising the possibility that loss of α-Cat may precipitate JNK activation through Rho1.

The loss of α-Cat leads to a corresponding decline of DEcad and Arm from AJs. Quantification of CCC proteins in the late stage embryonic epidermis of zygotic *α-Cat* null mutants suggested that AJs can be retained and support normal tissue architecture when CCC levels are reduced to less than 10% [Sarpal et al., 2012]. As heterozygous *α-Cat* animals are normal, we anticipate that reduction of α-Cat to somewhere between 50% and 10% of wild-type levels will cross a threshold that will increase Yki activity without activating JNK signaling sufficiently to cause cell death. How low levels of α-Cat and DEcad cause an activation of Yki remains to be explored. For example, reducing α-Cat could compromise the interactions between the AJ protein Echinoid and Salvadore, a Hippo binding partner important for normal Hippo activity [Yue et al., 2012], or compromise the interactions between Crumbs and Expanded that could deregulate Hippo signaling [Robinson et al., 2010; Ling et al., 2010; Chen et al., 2010]. Loss of α-Cat could also affect actin polymerization as mature AJs suppress actin polymerization [Drees et al., 2005; Sarpal et al., 2012; Maitre and Heisenberg, 2013], whereas enhanced actin polymerization is a known activator of Yki [Fernandez et al., 2011; Sansores-Garcia et al., 2011; Karaman and Halder, 2017]. Reducing α-Cat could therefore directly promote actin polymerization prior to an overt defect in cell adhesion, and consequently stimulate Yki activity and tissue growth. Finally, loss of α-Cat may affect Yki not through the Hippo pathway as was documented for αE-catenin in mammalian keratinocytes [Li et al., 2016].

The mechanosensitive interactions between Jub and α-Cat are thought to transmit tissue tension into growth regulatory signals. Cytoskeletal tension enhances recruitment of Jub to junctional α-Cat. As Jub forms a complex with Wts, Wts would be sequestered to AJs where it does not phosphorylate Yki, activating it [Rauskolb et al., 2014; Sun et al., 2015; Pan et al., 2016; Ibar et al, 2018]. Supporting this model, we observed Jub and Yki-dependent tissue overgrowth that correlates with an enhanced recruitment of Jub to AJs. In particular, our data suggest that the M1 domain acts as a gatekeeper for Jub recruitment. In the absence of M1, junctional Jub levels become strikingly high, suggesting that M region mechanosensing has become ineffective in limiting Jub recruitment, causing Yki activation and overgrowth [see also Alegot et al., 2019].

Tension is thought to cause a conformational change in the M region and an unfurling of the M1 domain exposing a Vinc binding site [Yonemura et al., 2010; Ishiyama et al., 2013; Yao et al., 2014; Kim et al., 2015; Barrick et al., 2018]. However, Vinc is not required for limiting Jub recruitment indicating that either a second unknown binding partner of M1 or possibly an intramolecular interaction between M1 and the Jub binding site in the N domain [Marie et al., 2003; Alegot et al., 2019; this work] controls Jub binding. Enhanced recruitment of Jub to AJs may also explain the tissue overgrowth resulting from α-Cat overexpression.

This Jub recruitment-dependent model of how mechanotransduction by the α-Cat M region regulates tissue growth does not explain all our observations and therefore needs to be extended to incorporate additional mechanisms of how AJ can control tissue growth. First, this model cannot explain the Yki-dependent overgrowth precipitated by low α-Cat levels, and corresponding low Jub levels at AJs, which we have discussed above. Second, we observed that two fusion proteins between DEcad and α-Cat, one containing both full-length proteins (DEcad::αCat) and one lacking the N-domain of α-Cat (DEcadΔβ::αCatΔNT-N), can both support normal growth in α-Cat KD tissue but only DEcad::αCat restored normal junctional Jub levels whereas DEcadΔβ::αCatΔNT-N did not. Third, expression of α-CatR in α-Cat KD tissue restores α-Cat to approximately 63% of normal levels. However, Jub increases to 119% without a noticeable increase in tissue size. One possibility is that mechanical force distributed over fewer α-Cat molecules enhances the mechanosensory response of individual α-Cat molecules in a non-linear manner resulting in higher Jub recruitment. Evidence for such a mechanism was recently reported for the recruitment of Vinc to AJs in the Drosophila germband [Kale et al., 2018], and may be similar for Jub recruitment to AJ in that tissue [Razzell et al., 2018]. Fourth, removing the entire M region results in an α-Cat protein that can support normal wing development, implying that M region mechanosensing is not an essential aspect of regulating tissue growth. In light of the results with α-CatR-ΔM1, this can only be explained by assuming that removing M2 and M3 in addition to M1 has a compensatory effect. Whereas α-CatR and α-CatR-ΔM expression in α-Cat depleted PC tissue enhances Jub recruitment above AC control levels, loss of M2 or M3 causes Jub levels to remain below AC levels suggesting that these two domains somehow normally support α-Cat ability to recruit Jub. Collectively, our findings suggest that M region mechanosensing contributes to Jub recruitment and Hippo/Yki pathway regulation but that the actual mechanisms involved have considerable complexity and require further analysis to be resolved.

Increased levels of Jub were also observed in response to disrupting the mechanosensory properties of the α-Cat ABD. Disruption of the α1-helix of ABD did not only enhance F-actin binding in vitro [Ishiyama et al., 2018] but also stabilized the CCC at AJs in the wing epithelium as suggested by our observations. A corresponding increase in Jub levels at AJs could account for the persistent overgrowth observed in α-Cat KD tissue expressing α1-helix compromised α-Cat (α-CatR-H1). Thus, the mechanosensory properties of the M region and the α-Cat ABD are both important for regulating Jub recruitment to AJs and growth regulation through the Hippo/Yki pathway. As changes in tissue tension are thought to modulate the Hippo/Yki pathway through junctional recruitment of Jub [Rauskolb et al., 2014; Eder et al., 2017] we asked whether α-Cat mutants that compromise mechanosensing cause changes in tissue tension. We did not observe such changes as assessed through studying the initial recoil velocities of cell vertices after laser ablation of the connecting junction. These findings suggest that α-Cat operates as a crucial mechanosensor to regulate tissue growth.

In summary, we conclude that α-Cat uses multiple mechanisms to act as an important regulator of tissue growth in the Drosophila wing disc epithelium. It is doing so at least in part by operating as a mechanotransducer, engaging both M region and ABD mechanosensing, to relay cytoskeletal tension into growth regulatory signals. One of these mechanisms involves the recruitment of Jub to the N domain of α-Cat that can be modulated by both mechanosensing mechanisms. However, α-Cat also engages mechanisms that are independent of the junctional recruitment of Jub to control tissue growth, which remain to be uncovered.

## Materials and Methods

### Drosophila stocks

The fly stocks used were as follows: *Ubi-α-Cat Tub-GAL80 FRT40A (Sarpal et al., 2012), hsFLP FRT40A;α-Cat^1^ da-GAL4 UAS-mCD8::GFP/TM6B (Sarpal et al., 2012), UAS-α-Cat-RNAi(1)/TM6B* (this work), *en-Gal4 UAS-RFP/CyO* (Bloomington Drosophila Stock Center (BDSC) 30557), *UAS-p35* (Hay et al., 1994), *puc^E697^-lacZ/TM6B* (Martin-Blanco et al., 1997), *UAS-shg-RNAi* (Vienna Drosophila Resource Center (VDRC) 27081), *omb-Gal4), ex-lacZ/CyO* (Bloomington Drosophila stock center), *UAS-α-Cat-RNAi(2)* (TRiP HMS00317, Harvard Medical School), *UAS-jub-RNAi* (TRiP line HMS00714, Harvard Medical School), *UAS-ex-RNAi* (TRiP line HMS00874, Harvard Medical School), *UAS-crb-RNAi* (TRiP line JF02777, Harvard Medical School), *UAS-ft-RNAi* (TRiP line HMS00932, Harvard Medical School), *yki^B5^/CyO* (Huang et al., 2005), *p120^308^* (Myster et al., 2003), *Jub::GFP* (Sabino et al., 2011), and *Vinc* (Klapholz et al., 2015).

### Clonal analysis

*α-Cat^1^* MARCM clones in imaginal discs and follicle cells were generated as described (Sarpal et al., 2012). Briefly, to induce clones in wing discs, embryos were collected at 25°C for 24 hours, larvae were heat shocked twice at 37°C for 2 hours at 48 hours after egg laying (AEL), and again at 72 hours AEL. Larvae were collected at 96 hrs AEL and dissected. Clones were induced in larvae of the following genotype:

- *hs-FLP FRT40A/tub-Gal80 Ubi-α-Cat FRT40A; Act5c-Gal4 /da-Gal4 UAS-mCD8::GFP αCat^1^*
- *hs-FLP FRT40A/tub-Gal80 Ubi-α-Cat FRT40A; Act5c-Gal4 α-Cat^1^/da-Gal4 UAS-mCD8::GFP αCat^1^* and
- *hs-FLP FRT40A/tub-Gal80 Ubi-α-Cat FRT40A; Act5c-Gal4 UAS-p35 α-Cat^1^/da-Gal4 UAS-mCD8::GFP α-Cat^1^*.

To induce follicle-cell clones, flies were grown at 25°C. Pupae were heat shocked at 37°C for 2 hours each on two consecutive days and then shifted to 29°C until the pharate adult stage, and then shifted back to 25°C. Freshly eclosed females were collected and moved to yeasted vials with a couple of males to induce oogenesis and then dissected after 2 days. Clones were induced in flies of the following genotype:

- *hs-FLP FRT40A/tub-Gal80 Ubi-α-Cat FRT40A; Act5c-Gal4 α-Cat^1^/da-Gal4 UAS-mCD8::GFP αCat^1^* and
- *hs-FLP FRT40A/tub-Gal80 Ubi-α-Cat FRT40A; Act5c-Gal4 α-Cat^1^ UAS-X/da-Gal4 UAS-mCD8::GFP αCat^1^*.
*UAS-X* refers to *UAS-α-CatR* or *UAS-α-CatΔM*.

### Generation of *Drosophila* transgenes

To generate the *UASp-α-Cat* constructs, full-length α-Cat cDNA (Oda and Tsukita 1999; 2751 nucleotides) was cloned into Gateway pENTR™/D-TOPO™ entry vector (K240020, Thermo Fischer Scientific) digested with *NotI* and AscI, using 3-part Gibson assembly reaction (New England Biolabs). *αCat* cDNAs carrying various mutations (Suppl. Table S2) were cloned using 2-part or 3-part Gibson assembly reactions with *UASp-α-Cat* in pENTR™/D-TOPO™ as the backbone digested with restriction enzymes. DNA fragments carrying the mutations were synthesized by Thermo Fischer. The Gateway^®^ LR^®^ Clonase Enzyme mix was used to clone all entry vector constructs into *pPWH-attB* (pUASP-Gateway Cassette with C-terminal 3xHA tag, #1102, Drosophila Genomics Resource Center modified by insertion of an *attB* recombination site (Groth et al., 2004) at a NSiI restriction site. Transgenic animals were produced by Best Gene Inc., by using flies carrying the *attP2* recombination site (Groth et al., 2004).

*UAS-α-Cat-RNAi(1)* construct was designed by selecting a 21-nt target sequence 5’-GGTTAAAGAATTTATGTTAAA-3’ in the *α-Cat* transcript based on the algorithm by Vert et al., 2006. The 71 basepair oligonucleotide sequence 5’-ctagcagtGGTTAAAGAATTT-ATGTTAAAtagttatattcaagcataTTTAACATAAATTCTTTAACCgcg-3’ with overhangs for the restriction enzymes Nhe1 and EcoR1 was produced using the shRNA protocol of Ni et al., 2011. The annealed oligonucleotides were cloned into the VALIUM20 vector (TRiP functional genomics resources, Harvard Medical School). To render α-CatR and derivative constructs resistant to the shRNAs *α-Cat-RNAi(1) and α-Cat-RNAi(2)* we made the following modifications: We changed the 5’-ATGTTAAAA-3’ sequence at the start of the *α-Cat* coding region, which is targeted by *UAS-α-Cat-RNAi(1)*, to 5’-ATGCT GAA G-3’, and we changed 5’-GCAGCATCGATATTGACTGTT-3’, which encodes sequences in the M2 domain and is targeted by *UAS-α-Cat-RNAi(2)* [TriP line HMS00317] to 5’-GCCGCCTCCATCCTGACCGTG-3’. The DEcadΔβ::αCat fusion protein was engineered similar to methods described in Sarpal et al., 2012. To remove the Arm binding domain in DEcadΔβ amino acids 1445-AYEGDGNSDGSLSSLASCTDD-1466 were deleted from the DEcad.

### Immunohistochemistry

Crosses using *α-CatRNAi(2)* were grown at 25°C. For *α-CatRNAi(1)*, embryos were collected for 24 hrs at 25°C and kept at 25°C until 48 hrs after egg laying (AEL), when they were moved to 29°C to enhance RNAi KD. Wing imaginal discs from late third instar larvae dissected in cold Phosphate buffer saline (PBS), and fixed for 20 min in 4% paraformaldehyde in PBS at room temperature, then washed 3 times in 0.1% PBS-Triton X-100 followed by 30 minutes of incubation in 0.3% PBS-Triton X-100. Tissue was incubated with 5% goat serum in 0.1% PBS-Triton X-100 and then processed for antibody staining. Primary antibodies used were: guinea pig polyclonal antibody (pAb) anti-α-Catenin (p121; 1:1000, Sarpal R. et al., 2012), rat monoclonal antibody (mAb) anti-HA (3F10, 1:500, Sigma), mouse mAb anti-Arm (N2-7A1, 1:50, Developmental Studies Hybridoma Bank [DSHB]), mouse mAB anti-β-Galactosidase antibody (Z378A, 1:500, Promega), pAb rabbit anti-active JNK (V7931, 1:100, Promega) and pAb rat anti-Crb antibody (F3, 1:500, Pellikka M. et al., 2002). Fluorescent secondary antibodies were used at a dilution of 1:400 (Jackson Immuno Research Laboratories and Invitrogen). Larval tissues were further incubated with DAPI (1:1000, Molecular Probes) or Acti-stain Phalloidin (PHDG1-A, 1:50, Cytoskeleton Inc.), followed by washes in 0.1% PBS-Triton X-100. Tissues were incubated in Vectashield^®^ antifade mounting medium (H-1000, Vector Laboratories) overnight. Wing discs were mounted apical side up in Vectashield, with one layer of Scotch double-sided tape on both short edges of the coverslips (VWR 22×40 mm, No. 1.5) as spacers to prevent excessive compression of the discs, as described by Spratford & Kumar, 2014. For experiments in Figures 8G and H, male larvae of the genotypes, *en-Gal4 UAS-RFP/jub::GFP* and *Vinc; jub::GFP/+* were fixed and processed together in the same Eppendorf tube and mounted on the same slide.

### RNAi KD, overexpression, and genetic interaction experiments in the adult wing

All adult wing crosses were set up using the *omb-Gal4* driver. Progeny were collected for 24 hrs at 25°C, and kept at 25°C until their emergence from pupae. Female progeny were collected for 48 hrs after hatching. Wings were then removed using forceps, washed in 0.1% PBS-Triton, and mounted in 50% Glycerol.

### Image acquisition, processing and quantification

Confocal images of wing discs were acquired on a Leica TCS SP8 scanning confocal microscope using HC PL APO CS2 20X/0.75 or HC PL APO CS2 40X/1.30 oil immersion objectives. Z-stacks were acquired to capture the full disc, and subsequent processing and maximal projections were made using the “Z-stack” function of Fiji (ImageJ, Akiyama and Gibson 2015). Total disc sizes were quantified by converting fluorescent to binary images using the Fiji threshold tool. Size measurements were made from outlines of total wing disc. Adult wing images were acquired on a Zeiss Axiophot microscope using the bright field mode, with the 5x lens (NA 0.15) objective. Subsequent processing and wing size quantification was performed using the “measure” function of Fiji. The wing size was quantified by converting the image to binary using the Fiji threshold tool, then using the Magic Wand tool to outline the wing, and the “Measure Area” tool to produce a pixel number for the area of the wing. Scale bars were generated using the “Analyse” tool in Fiji. Images were assembled in Adobe Photoshop and Adobe Illustrator. All graphs and statistical analyses were generated using Prism (Graph Pad) software. We used unpaired two-tailed *t* tests to determine p-values when samples passed the normality test, and we used nonparametric two-sample Kolmogorov-Smirnov (KS) test otherwise.

### Arm and Jub protein quantification

We used Scientific Image Segmentation and Analysis [SIESTA; Fernandez-Gonzalez and Zallen, 2011; Leung and Fernandez-Gonzalez, 2015] to automatically identify cell outlines on confocal stacks of wing discs to determine Arm and Jub levels. 1-pixel-wide lines (60 nm/pixel) were selected to obtain the mean pixel intensity per cell. Using SIESTA, ~200 cells were selected above and below the dorso-ventral compartment boundary in both the anterior and posterior compartments. To eliminate background from measurements of protein levels, we subtracted the values of protein level measurements for the center of the cell from protein level measurements for the cell perimeter. In case of *UAS-α-Cat-ΔM2, UAS-α-CatR-ΔβH* and *UAS-α-CatR-H1-ΔβH*, anti-HA staining was used as a reference to identify cell outlines in the posterior compartment. For every larval wing disc, average protein measurements for the anterior compartment thus calculated were used to normalize the protein measurements per cell selected from the posterior compartment. Average protein measurements were then plotted as a percentage. Statistical significance was calculated using nonparametric two-sample Matt Whitney test in Prism 7 (GraphPad Software).

### Live imaging of wing discs and laser ablation

Wing discs from wandering 3rd instar larvae were dissected and mounted in a solution containing 2% heat inactivated FBS, 10μg/ml insulin, and 1x Penicillin-Streptomyocin in Schneider’s medium. The discs were mounted between a cover glass and an oxygen permeable membrane. A Revolution XD spinning disk confocal (Andor) with a 100× oil-immersion lens (NA 1.40; Olympus), iXon Ultra 897 camera (Andor) and Metamorph (Molecular Devices) as the image acquisition software were used for imaging the laser ablation experiments. 16-bit Z-stacks of 7 slices each, at 0.2 μm per step, taken every 3 seconds for 2.5 minutes were used. Cells which were within the pouch, several cell diameters away from the AP and DV boundaries, roughly symmetrical in shape and without obvious polarization were chosen. Junctions of disc pouch cells located immediately below peripodial cell junctions or nuclei were avoided. A pulsed Micropoint N2 laser (Andor) tuned to 365 nm was used to apply ten laser pulses to a spot in the center of a single cell junction, with an image taken immediately before and after ablation (with 1.72s between images). Measurement of recoil velocity was performed by manually tracking the positions of tricellular vertices which were connected by the ablated edge using SIESTA, and the velocity of retraction was calculated using a MATLAB script provided by the Fernandez-Gonzalez lab.

### Whole-animal rescue experiments

Whole animal rescue experiments for *α-CatR, α-CatR-ΔM, DEcadΔβ::αCatΔN* were carried out as described [Desai et al., 2013]. Statistical significance was determined with a non-parametric Kolmogorov-Smironov test using Prism 7 (GraphPad software).

## Acknowledgements

We thank Alexa Chioran, Meggie Cao, Arun Shipstone, Limin Wang, and Marylin Rowswell for technical assistance, and Gayaanan Jeyanathan, Sergio Simoes, Rodrigo Fernandez-Gonzalez, and Arun Sarpal for helpful discussions. We thank Rodrigo Fernandez-Gonzalez for the use of his laser ablation setup and him and his students Jessica Yu, Anna Kobb and Teresa Zulueta-Coarasa for help with data analysis. Sergio Simoes and Dorothea Godt read the manuscript critically and made helpful suggestions. We gratefully acknowledge the use of the imaging facility of the Department of Cell and Systems Biology led by Henry Hong. This work was supported by a grant from the Canada First Research Excellence Fund (Government of Canada) through the Medicine-by-Design program at the University of Toronto, and by a Project Grant of the Canadian Institutes for Health Research. UT is a Canada Research Chair (Tier 1) for Epithelial Polarity and Development.

## Author’s contributions

R.S., V.Y., L.K., and L.S designed, carried out, and analysed experiments. U.T. conceptualized the project, raised funds, supervised R.S., V.Y., L.K., and L.S, and wrote the paper.

