## Supplemental Materials for "Role of α-Catenin and its mechanosensing properties in the regulation of Hippo/YAP-dependent tissue growth"

**Manuscript date March 4, 2019**

Character count (no spaces) (excluding Materials and Methods, References, and Suppl. Materials): 48,812

### Supplemental Materials

**Figure S1: Over-expression of α-Cat causes overgrowth in a sensitized background.**
**(A-B)** Depletion of Crb (A) or Ex (B) does not cause a significant enlargement of larval wing disc. However, overexpression of α-Cat in conjunction with a KD of (A) Crb and (B) Ex show synergistic overgrowth phenotypes (quantification shown in Figure 3D). Scale bars, 100 μm.

**Figure S2: Overgrowth resulting from over-expression of DEcad::α-Cat is suppressed by downregulation of Yki and Jub.**
**(A)** Overexpression of DEcad::α-Cat causes an enlargement of the adult wing, which can be suppressed by introducing one mutant copy of *yki* or expression of Jub ShRNA. **(B)** Quantification of adult wing areas. Two-tailed, unpaired t-test used to determine statistical significance. ****(P<0.0001).

**Figure S3: Levels of α-Cat are reduced compared to endogenous protein in α-Cat mutant cells expressing α-CatR.**
**(A,B)** Late 3rd larval instar wing discs of indicated genotypes labeled with DAPI and posterior compartment marked by RFP. Close-up images to the right show wing pouch area on both sides of the anterior-posterior compartment boundary. α-Cat was depleted in PC with *α-CatRNAi(2)*
**(C)** Comparison of relative fluorescent intensities between anterior compartment (AC) and posterior compartment (PC) for α-Cat. AC values were normalized to 100%. N=300-400 cells from two wing discs. Mann Whitney test was used to determine statistical significance. ****(P<0.0001)
**(D)** Western blot analysis of protein levels of α-CatR and α-Cat using anti-HA antibody. β-tubulin is used as the loading control.

**Figure S4: Deletion of the entire M-region has a minor impact on follicular epithelium integrity and whole animal survival.**
**(A)** Follicular epithelium clones positively marked with GFP in indicated genotypes. *α-Cat* mutant cells show strongly reduced levels of Arm and display cytoplasmic aggregates of α-Spectrin.

Expression of α-CatR and α-CatR-ΔM in *α-Cat* mutant cells substantially rescues this defect. Scale bars 20 μm.

**(B)** Comparison of rescue activities of α-CatR and α-CatR-ΔM when expressed in *α-Cat* mutant cells in the follicular epithelium. Data are presented as mean±s.e.m.

**(C)** Whole animal survival plot showing average and total range of rescue activity of α-CatR, α-CatR-ΔM and DEcadΔβ::αCatΔNT-N when expressed in *α-Cat* zygotic mutant embryos. Data are presented as mean±s.d. A score of 0 was given to *α-Cat*$^{1}$ zygotic null mutant embryos which frequently displayed defects in head morphogenesis ('head open' phenotype). Embryos that displayed an enhancement of *α-Cat*$^{1}$ phenotype were given the following scores: (−2) embryonic lethal with both head open and a dorsal open phenotype indicating a failure of dorsal closure; (−1) embryonic lethal with both the head open defect and a hole in the dorsal epidermis indicating incomplete closure. For the rest of the animals that displayed rescue of the *α-Cat*$^{1}$ phenotype following scoring criteria were used: (1) embryonic lethal with weak head defects ('abnormal head'); (2) embryonic lethal with normal head; (3) lethal at first larval instar; (4) lethal at second larval instar; (5) lethal at third larval instar; (6) early pupa lethal; (7) late pupa lethal; (8) adult.

**Figure S5: Expression of α-Cat constructs in wing imaginal.**

Late 3$^{rd}$ larval instar wing discs of indicated genotypes were labeled with HA to detect the transgenic protein and Arm.

**Table S1: List of Genotypes for all panels and figures.**

**Table S2: α-Cat constructs used in this study.**

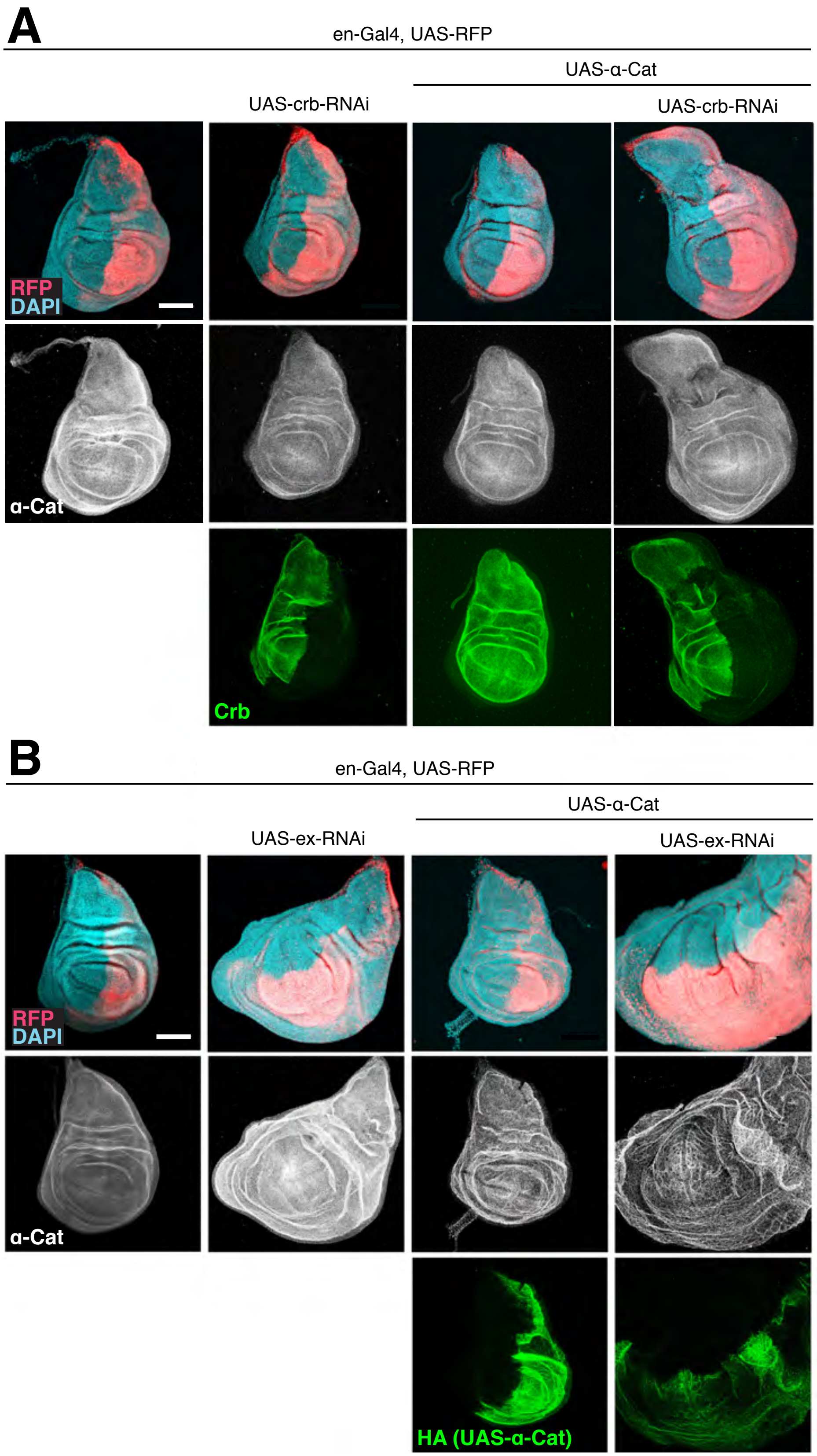


Sarpal et al., Figure S1

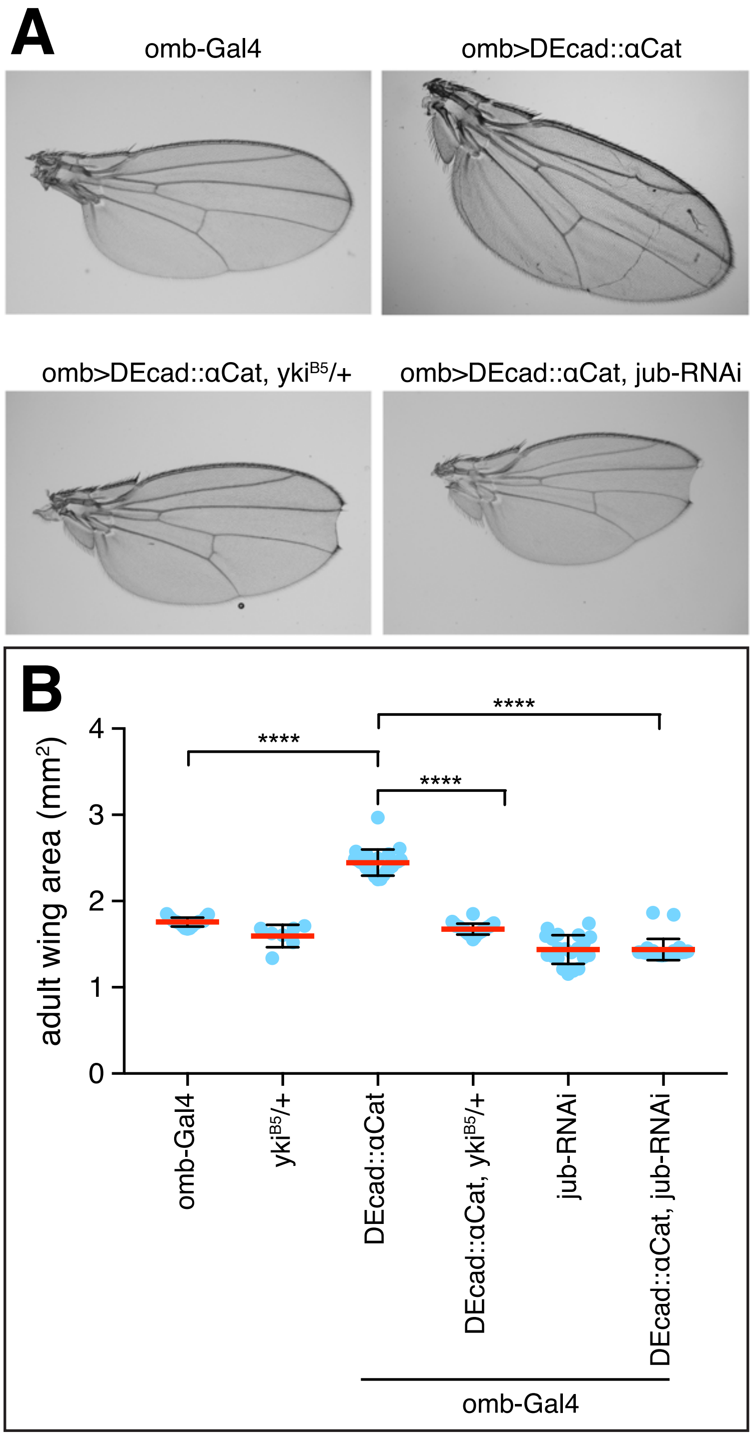


Sarpal et al., Figure S2

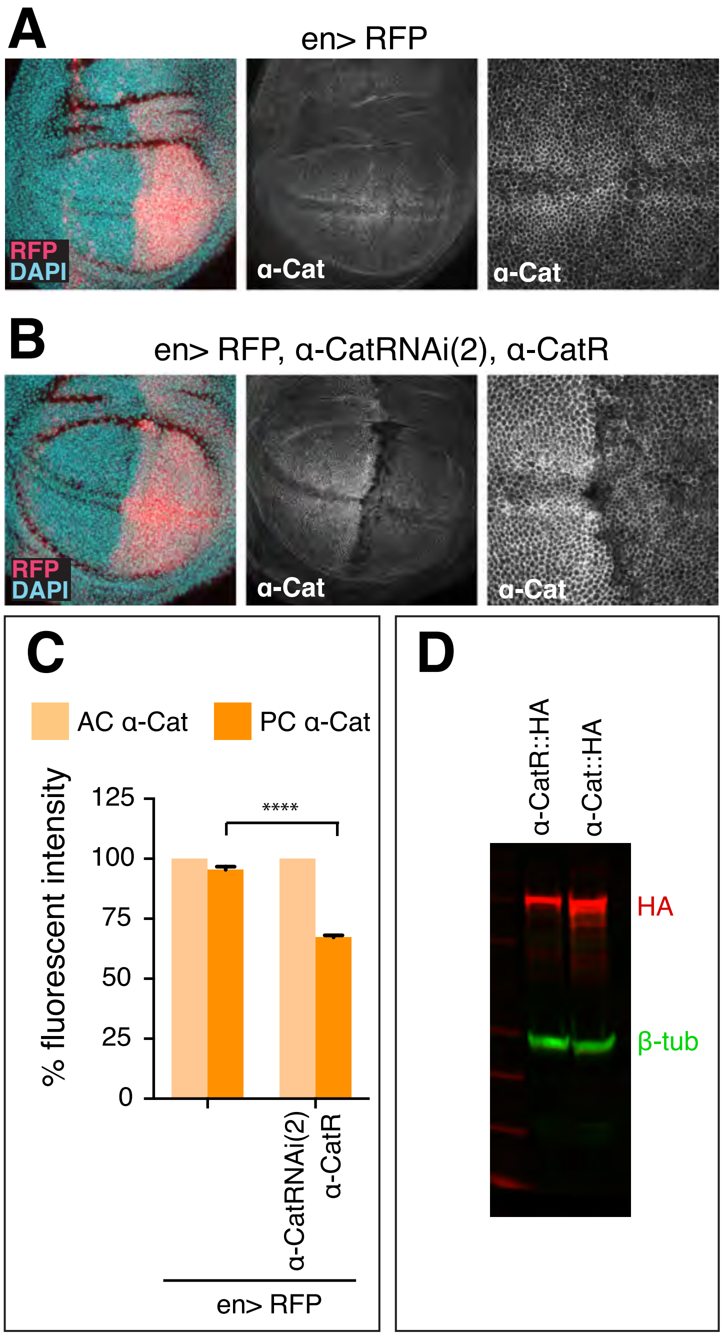


Sarpal et al., Figure S3

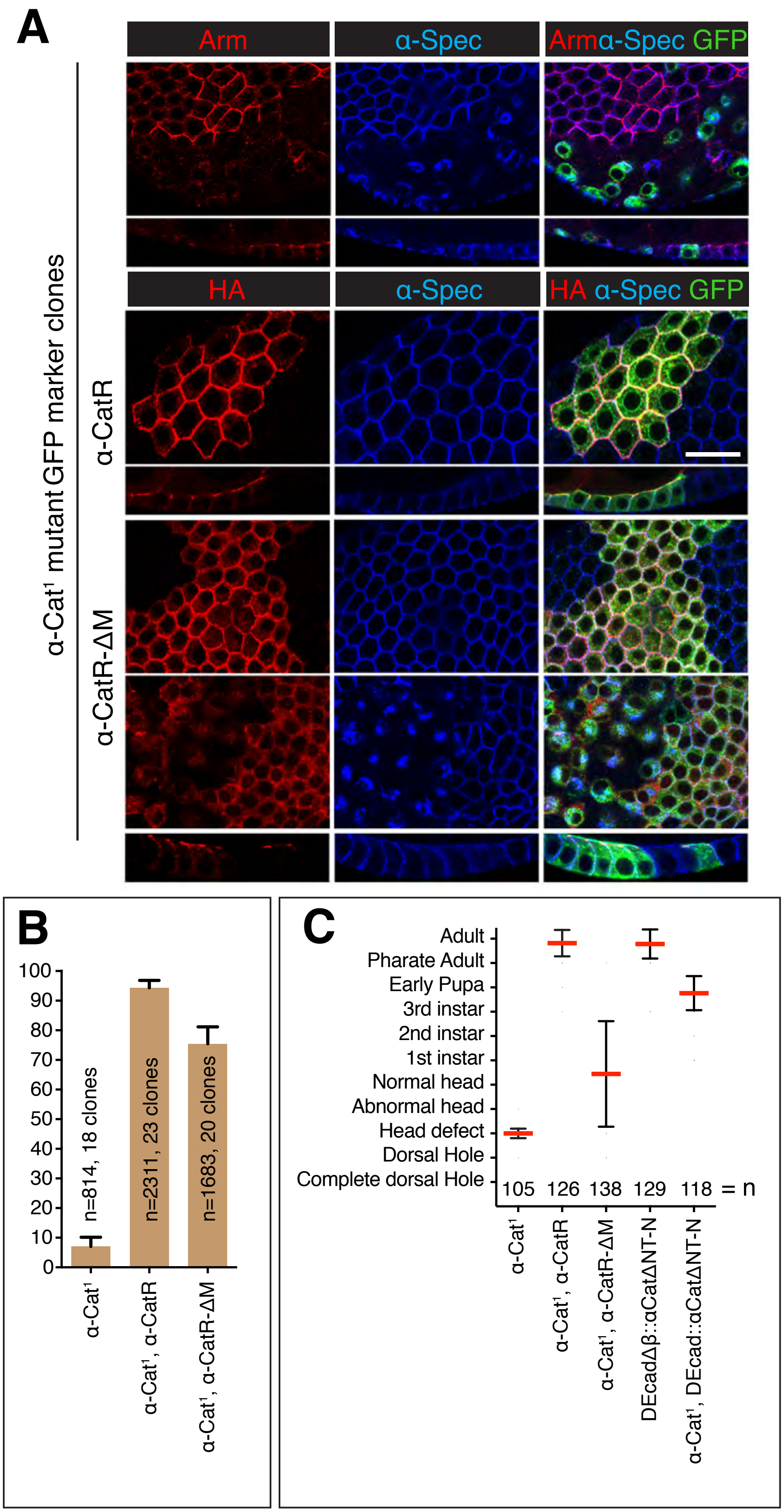


Sarpal et al., Figure S4

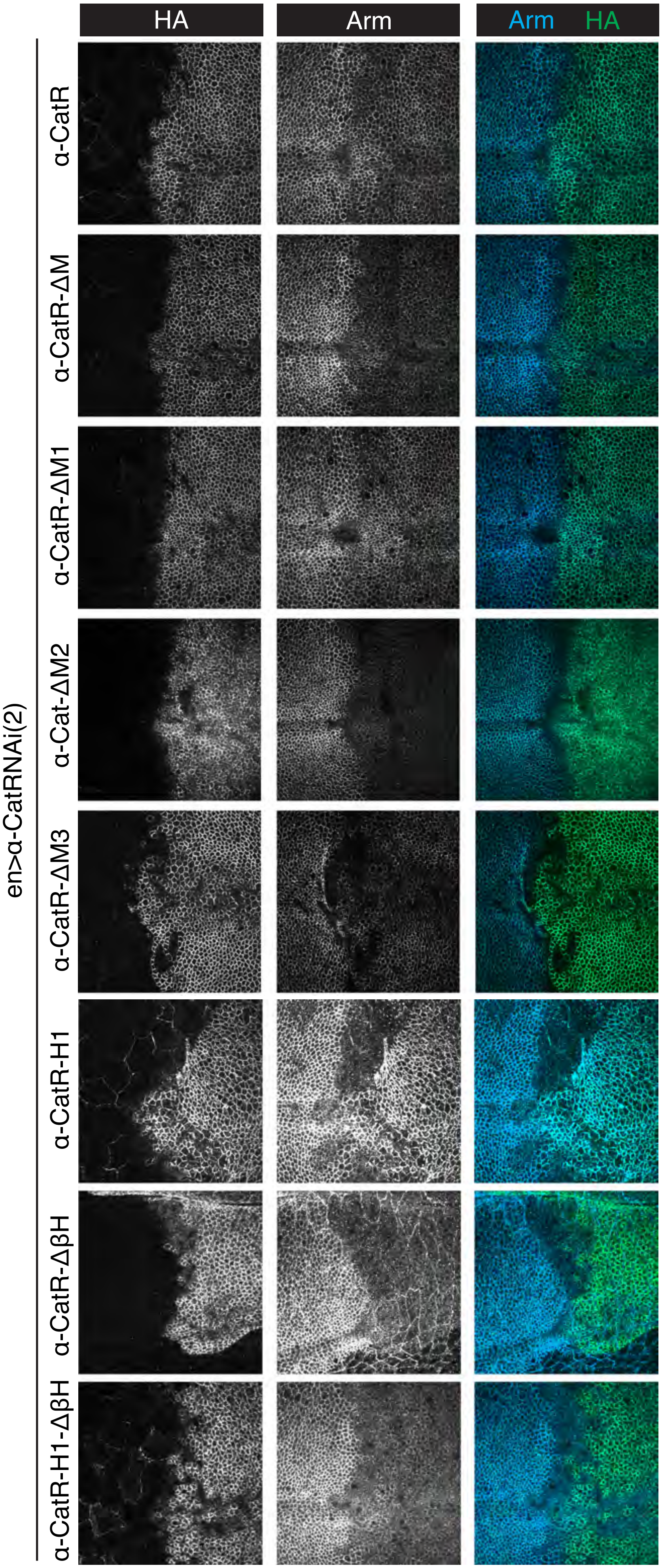


Sarpal et al.,
Figure S5

### Table S1: List of Genotypes for all panels and figures.

**Figure 1**
**B**:
Gal80, ubi-αcat, FRT40A/hsFLP ; Act-Gal4, α-Cat$^1$/Da-Gal4,UAS-GFP, α-Cat$^1$
Gal80, ubi-αcat, FRT40A/hsFLP; Act-Gal4, α-Cat$^1$, UAS-P35/Da-Gal4, UAS-GFP, α-Cat$^1$

**C**:
en-Gal4, UAS-RFP/+ ; UAS-α-Cat-RNAi$^1$/ +
en-Gal4, UAS-RFP/+ ; UAS-α-Cat-RNAi$^1$, α-Cat$^1$/+

**D**: Total wing disc measurements of the following genotypes are shown in the graph:

en-Gal4, UAS-RFP/+
en-Gal4, UAS-RFP/+; UAS-α-Cat-RNAi$^1$/ +
en-Gal4, UAS-RFP/+; UAS-α-Cat-RNAi$^1$/ UAS-P35
en-Gal4, UAS-RFP/+; UAS-α-Cat-RNAi$^1$, α-Cat$^1$/+
en-Gal4, UAS-RFP/+; UAS-α-Cat-RNAi$^1$, α-Cat$^1$/α-CatR
en-Gal4, UAS-RFP/+; UAS-α-Cat-RNAi$^2$/+
en-Gal4, UAS-RFP/+; UAS-α-Cat-RNAi$^2$/α-CatR

**E**:
en-Gal4, UAS-RFP/+ ; UAS-α-Cat-RNAi$^1$/UAS-P35
en-Gal4, UAS-RFP/+ ; UAS-α-Cat-RNAi$^1$, α-Cat$^1$/UAS-P35

**Figure 2**
**A:**
en-Gal4, UAS-RFP/Puc$^{E697}$-lacZ; UAS-α-Cat-RNAi$^1$/+
en-Gal4, UAS-RFP/Puc$^{E697}$-lacZ; UAS-α-Cat-RNAi$^1$/UAS-P35

**C:**
en-Gal4, UAS-RFP, ex-lacZ/+; UAS-α-Cat-RNAi$^1$/+
en-Gal4, UAS-RFP, ex-lacZ/+; UAS-α-Cat-RNAi$^1$, UAS-P35/+
en-Gal4, UAS-RFP/+; yki$^{B5}$/+
en-Gal4, UAS-RFP/yki$^{B5}$; UAS-α-Cat-RNAi$^1$/+

**D:** Total wing disc measurements of following genotypes are shown in the graph:
en-Gal4, UAS-RFP/+
en-Gal4, UAS-RFP/+; yki$^{B5}$/+
en-Gal4, UAS-RFP/+; UAS-α-Cat-RNAi$^1$/ +
en-Gal4, UAS-RFP/yki$^{B5}$; UAS-α-Cat-RNAi$^1$/+

**Figure 3**
**A:**
omb-Gal4/+
omb-Gal4/+; UAS-α-Cat-RNAi$^1$/+
omb-Gal4/+; UAS-α-Cat
yki$^{B5}$/+
omb-Gal4/+; yki$^{B5}$/+; UAS-α-Cat/+

**B:**
en-Gal4, UAS-RFP/+
en-Gal4, UAS-RFP/+; +/+; UAS-ft-RNAi/+

en-Gal4, UAS-RFP/+; UAS-α-Cat/+
en-Gal4, UAS-RFP, ex-lacZ/+; UAS-ft-RNAi/ UAS-α-Cat
en-Gal4, UAS-RFP/+; $yki^{B5}$/+; UAS-α-Cat/UAS-ft-RNAi

**C:** Adult wing measurements of following genotypes are shown in the graph:
omb-Gal4/+
$yki^{B5}$/+
omb-Gal4/+; UAS-α-Cat/+
omb-Gal4/+; $yki^{B5}$/+; UAS-α-Cat/+

**D:** Total wing disc measurements of following genotypes are shown in the graph:
en-Gal4, UAS-RFP/+
en-Gal4, UAS-RFP/+; +/+; UAS-ft-RNAi/+
en-Gal4, UAS-RFP, ex-lacZ/+; UAS-ft-RNAi/ UAS-α-Cat
en-Gal4, UAS-RFP/+; +/+; UAS-crb-RNAi/+
en-Gal4, UAS-RFP/+; +/+; UAS-crb-RNAi/ UAS-α-Cat
en-Gal4, UAS-RFP/+; +/+; UAS-ex-RNAi/+
en-Gal4, UAS-RFP/+; +/+; UAS-ex-RNAi/ UAS-α-Cat

**Figure 4**
**A:**
en-Gal4, UAS-RFP, ex-lacZ/+;
en-Gal4, UAS-RFP, ex-lacZ/+; +/+; UAS-DEcad-RNAi/+
en-Gal4, UAS-RFP, ex-lacZ/+; +/+; UAS-DEcad-RNAi/UAS-P35

**B:**
en-Gal4, UAS-RFP/$Puc^{E697}$-lacZ
en-Gal4, UAS-RFP/$Puc^{E697}$-lacZ; UAS-DEcad-RNAi/+
en-Gal4, UAS-RFP, ex-lacZ/+; +/+; UAS-DEcad-RNAi/UAS-P35

**C:** Total wing disc measurements of following genotypes are shown in the graph:
en-Gal4, UAS-RFP, ex-lacZ/+
en-Gal4, UAS-RFP, ex-lacZ/+; +/+; UAS-DEcad-RNAi/+
en-Gal4, UAS-RFP, ex-lacZ/+; +/+; UAS-DEcad-RNAi/UAS-P35

**D:** Total adult wing measurements of following genotypes are shown in the graph:
p120/p120

**Figure 5**
omb-Gal4/+
omb-Gal4/+; UAS-α-CatR/+
omb-Gal4/+; UAS-α-CatFL/+
omb-Gal4/+; UAS-α-CatR-ΔM/+
omb-Gal4/+; UAS-α-CatR-ΔM1/+
omb-Gal4/+; UAS-α-CatΔM2/+
omb-Gal4/+; UAS-α-CatR-ΔM3/+
omb-Gal4/+; UAS-DEcad/+
omb-Gal4/+; UAS-DEcad:: α-Cat/+
omb-Gal4/+; UAS-DEcadΔ🚲:: α-CatΔN/+
omb-Gal4/+; UAS-α-CatR-H1/+
omb-Gal4/+; UAS-α-CatR-Δ🚲H/+
omb-Gal4/+; UAS-α-CatR-H1-Δ🚲H /+
omb-Gal4/+; UAS-α-CatR-3A
omb-Gal4/+; UAS-α-CatΔABD/+

**Figure 6**
**A:**
en-Gal4, UAS-RFP/+; Jub::GFP/+

**B:**
en-Gal4, UAS-RFP/+; Jub::GFP/+; UAS-α-Cat-RNAi$^{1}$/+

**C:**
en-Gal4, UAS-RFP/+; Jub::GFP/+; UAS-α-Cat-RNAi$^{1}$/UAS-DEcad::α-Cat

**D:**
en-Gal4, UAS-RFP/+; Jub::GFP/+; UAS-α-Cat-RNAi$^{1}$/UAS-DEcadΔ🚲::α-CatΔN

**E:** Total wing disc measurements of following genotypes are shown in the graph:
en-Gal4, UAS-RFP/+
en-Gal4, UAS-RFP/+; UAS-α-Cat-RNAi$^{1}$/+
en-Gal4, UAS-RFP/+; UAS-α-Cat-RNAi$^{1}$/UAS-DEcad:: α-Cat
en-Gal4, UAS-RFP/+; UAS-α-Cat-RNAi$^{1}$/UAS-DEcadΔ🚲::α-CatΔN

**Figure 7:**
**A:**
en-Gal4, UAS-RFP/+
en-Gal4, UAS-RFP/+; UAS-α-Cat-RNAi$^{2}$/+
en-Gal4, UAS-RFP/+; UAS-α-Cat-RNAi$^{2}$/ UAS-α-CatR
en-Gal4, UAS-RFP/+; UAS-α-Cat-RNAi$^{2}$/ UAS-α-CatR-ΔM
en-Gal4, UAS-RFP/+; UAS-α-Cat-RNAi$^{2}$/ UAS-α-CatR-ΔM1
en-Gal4, UAS-RFP/+; UAS-α-Cat-RNAi$^{2}$/ UAS-α-Cat-ΔM2
en-Gal4, UAS-RFP/+; UAS-α-Cat-RNAi$^{2}$/ UAS-α-CatR-ΔM3

**B:**
omb-Gal4/+
omb-Gal4/+; UAS-α-CatR-ΔM1
yki$^{B5}$/+
omb-Gal4/+; yki$^{B5}$/+; UAS-α-CatR-ΔM1
omb-Gal4/+; JubRNAi/+
omb-Gal4/+; JubRNAi/+; UAS-α-CatR-ΔM1/+

**C:** Total wing disc measurements of following genotypes are shown in the graph:
en-Gal4, UAS-RFP/+
en-Gal4, UAS-RFP/+; UAS-α-Cat-RNAi$^{2}$/+
en-Gal4, UAS-RFP/+; UAS-α-Cat-RNAi$^{2}$/ UAS-α-CatR-ΔM
en-Gal4, UAS-RFP/+; UAS-α-Cat-RNAi$^{2}$/ UAS-α-CatR-ΔM1
en-Gal4, UAS-RFP/+; UAS-α-Cat-RNAi$^{2}$/ UAS-α-Cat-ΔM2
en-Gal4, UAS-RFP/+; UAS-α-Cat-RNAi$^{2}$/ UAS-α-CatR-ΔM3

**D:** Total adult wing measurements of following genotypes are shown in the graph:
omb-Gal4/+
omb-Gal4/+; UAS-α-CatR-ΔM1/+
yki$^{B5}$/+
omb-Gal4/+; yki $^{B5}$/+; UAS-α-CatR-ΔM1/+
omb-Gal4/+; Jub-RNAi
omb-Gal4/+; JubRNAi/ UAS-α-CatR-ΔM1

**Figure 8:**

**A:**
en-Gal4, UAS-RFP/+; Jub::GFP/+; UAS-α-Cat-RNAi$^{2}$/ UAS-α-CatR

**B:**
en-Gal4, UAS-RFP/+; Jub::GFP/+; UAS-α-Cat-RNAi$^{2}$/ UAS-α-CatR-ΔM

**C:**
en-Gal4, UAS-RFP/+; Jub::GFP/+; UAS-α-Cat-RNAi$^{2}$/ UAS-α-CatR-ΔM1

**D:**
en-Gal4, UAS-RFP/+; Jub::GFP/+; UAS-α-Cat-RNAi$^{2}$/ UAS-α-Cat-ΔM2

**E:**
en-Gal4, UAS-RFP/+; Jub::GFP/+; UAS-α-Cat-RNAi$^{2}$/ UAS-α-CatR-ΔM3

**F:**
Armadillo and Ajuba fluorescence intensity quantifications of genotypes listed in A,B,C,D,E and F above.

**G:**
en-Gal4, UAS-RFP/+; Jub::GFP/+
$vinc^{102.1}$/Y; Jub::GFP/+

**H:** Total wing disc measurements of following genotypes are shown in the graph:
en-Gal4, UAS-RFP/+
$vinc^{102.1}$/Y; Jub::GFP/+

**I:** Ajuba::GFP fluorescence intensity of following genotypes are shown in the graph:
en-Gal4, UAS-RFP/+
$vinc^{102.1}$/Y; Jub::GFP/+

**Figure 9:**
**A:**
en-Gal4, UAS-RFP/+; UAS-α-Cat-RNAi$^{2}$/ UAS-α-CatR
en-Gal4, UAS-RFP/+; UAS-α-Cat-RNAi$^{2}$/ UAS-α-CatR-H1
en-Gal4, UAS-RFP/+; UAS-α-Cat-RNAi$^{2}$/ UAS-α-CatR-Δ🚲H
en-Gal4, UAS-RFP/+; UAS-α-Cat-RNAi$^{2}$/ UAS-α-CatR-H1-Δ🚲H
en-Gal4, UAS-RFP/+; UAS-α-Cat-RNAi$^{1}$/ UAS-α-Cat3A
en-Gal4, UAS-RFP/+; UAS-α-Cat-RNAi$^{1}$/ UAS-α-CatΔABD

**B:** Total wing disc measurements of genotypes in A are shown in the graph:

**C:**
en-Gal4, UAS-RFP/+; Jub::GFP/+; UAS-α-Cat-RNAi$^{2}$/ UAS-α-CatR-H1

**D:**
en-Gal4, UAS-RFP/+; Jub::GFP/+; UAS-α-Cat-RNAi$^{2}$/ UAS-α-CatR-Δ🚲H

**E:**
en-Gal4, UAS-RFP/+; Jub::GFP/+; UAS-α-Cat-RNAi$^{2}$/ UAS-α-CatR-H1-Δ🚲H

**F:** Armadillo and Ajuba fluorescence intensity of following genotypes are shown in the graph:
en-Gal4, UAS-RFP/+; Jub::GFP/+; UAS-α-Cat-RNAi$^{2}$/ UAS-α-CatR
en-Gal4, UAS-RFP/+; Jub::GFP/+; UAS-α-Cat-RNAi$^{2}$/ UAS-α-CatR-H1
en-Gal4, UAS-RFP/+; Jub::GFP/+; UAS-α-Cat-RNAi$^{2}$/ UAS-α-CatR-Δ🚲H

en-Gal4, UAS-RFP/+; Jub::GFP/+; UAS-α-Cat-RNAi$^2$/ UAS-α-CatR-H1-Δ🚲H

**Figure 10:**
**A,B:**
en-Gal4, UAS-RFP/+; Jub::GFP/+; UAS-α-Cat-RNAi$^2$/ UAS-α-CatR
en-Gal4, UAS-RFP/+; Jub::GFP/+; UAS-α-Cat-RNAi$^2$/ UAS-α-CatR-ΔM
en-Gal4, UAS-RFP/+; Jub::GFP/+; UAS-α-Cat-RNAi$^2$/ UAS-α-CatR-ΔM1
en-Gal4, UAS-RFP/+; Jub::GFP/+; UAS-α-Cat-RNAi$^2$/ UAS-α-CatR-H1

**C:** Initial retraction velocity for genotypes in A are shown in the graph.

| Table S1 | | | | |
|---|---|---|---|---|
| **Constructs** | **Amino Acids (a-Catenin)** | **Gateway cloning** | **HA tag** | **Reference** |
| **α-Cat Full Length** | | | | |
| α-CatR | 1-917 | yes | yes | this work |
| α-Cat | | no | yes | Desai et al., 2013 |
| **DEcad-αCat Fusion Constructs** | | | | |
| DEcad | 0 | no | no | Sarpal et al., 2012 |
| DEcad::α-Cat | 1-917 | no | no | Sarpal et al., 2012 |
| DEcadΔß::α-CatΔNT-N | 281-917 | no | no | this work |
| **M domain Constructs** | | | | |
| α-CatR-ΔM | 1-292 / 635-917 | yes | yes | this work |
| α-CatR-ΔM1 | 1-292 / 399 -917 | yes | yes | this work |
| α-Cat-ΔM2 (α-Cat-ΔVH2N) | 1-399 / 509-917 | no | yes | Desai et al., 2012 |
| α-CatR-ΔM3 | 1-509 / 635-917 | yes | yes | this work |
| **ABD Constructs** | | | | |
| α-CatR-H1 | 683REAM > 683GSGS | yes | yes | this work |
| α-CatR-ΔßH | 1- 811/824-917 | yes | yes | Ishiyama et al., 2018 |
| α-CatR-H1-ΔßH | 1-811/824-917 + 683REAM > 683GSGS | yes | yes | this work |
| α-CatR-3A | L798A + I805A + V809A | yes | yes | Ishiyama et al., 2018 |
| α-CatR-ΔABD | 1-681 | yes | yes | this work |
